# Patient-derived IgG Amplifies Fcγ Receptor-Dependent Colonic Inflammation During Immune Checkpoint Blockade

**DOI:** 10.64898/2026.06.03.729692

**Authors:** Iryna Voloshyna, Yury Patskovsky, Sabina Sandigursky, Chaitra Sreenivasaiah, Erol C. Bayraktar, Ethan Tardio, Andrea V. Lopez, Stanzin Idga, Courtney Ng, Milad Ibrahim, Clara Goldberg, Anastasia Zhurova, Ryah Freih, Justin Mastroianni, Yuan Hao, Pamela Mishra, Alireza Khodadadi-Jamayran, Janice Mehnert, Gregg J. Silverman, Faisal Fa’ak, Iman Osman, Michelle Krogsgaard

**Affiliations:** Laura and Isaac Perlmutter Cancer Center, New York University Grossman School of Medicine, New York, NY 10016, USA; Department of Pathology, New York University Grossman School of Medicine, New York, NY, 10016, USA; Ronald O. Perelman Department of Dermatology, New York University Grossman School of Medicine, New York, NY, 10016, USA; Applied Bioinformatics Laboratories, New York University Langone Medical Center, New York, NY, 10016, USA; Department of Medicine, New York University Grossman School of Medicine, New York, NY, 10016, USA

## Abstract

Immune checkpoint inhibitor (ICI)-associated colitis limits effective cancer immunotherapy, yet host determinants of severe toxicity remain undefined. We investigated whether humoral immunity is associated with subsequent severe immune-related colitis (irC). In melanoma patients, baseline serum autoantibody (AAb) profiling identified a composite antigen signature - a signature-level association rather than validated functional specificities - associated with severe irC, marked by retained reactivity to tumor-associated antigens and relative depletion of antibodies recognizing immune- and mucosal-regulatory proteins. To assess functional relevance, we transferred polyclonal IgG from severe- or non-severe-irC patients into wild-type or humanized Fcγ receptor (hFcγR) mice treated with anti-PD-1 or anti-CTLA-4. IgG alone did not induce inflammation; however, severe- irC IgG amplified checkpoint-driven colonic inflammation in hFcγR mice, but not in wild-type mice, with checkpoint-specific remodeling of myeloid, lymphoid, and innate lymphoid compartments. These findings suggest that pretreatment humoral immunity conditions susceptibility to irC via the IgG-FcγR axis.

**Statement of Significance:** Pretreatment autoantibody signatures represent host humoral states associated with risk of severe immune checkpoint inhibitor-associated colitis. Patient-derived polyclonal IgG amplified checkpoint-driven intestinal inflammation through FcγR-dependent mechanisms without independently inducing disease, supporting the IgG-FcγR axis as a candidate pathway for mechanistic study and toxicity-risk stratification.

## INTRODUCTION

Immune checkpoint inhibitors (ICIs) have been a cornerstone of cancer immunotherapy for more than a decade, producing durable benefit across multiple malignancies [1]. Their therapeutic success, however, is limited by immune-related adverse events (irAEs), which can affect nearly any organ system and remain a major cause of treatment interruption and morbidity [2, 3]. Among these toxicities, immune-related colitis [4] is one of the most frequent and clinically consequential [5, 6]. Despite its clinical importance, the host factors that determine susceptibility to severe irC remain poorly defined.

Current models of irC are largely centered on modifications of cellular immunity triggered by checkpoint blockade, including impaired regulatory T cell (Treg)-function, loss of IL-10- and TGFβ-dependent immune restraint, and expansion of Th1-, Th17-, and cytotoxic effector programs [2, 6, 8–10]. Single-cell studies of melanoma-associated irC reinforced this model, demonstrating the involvement of ITGAE^hi^ and ITGB2^hi^ CD8 tissue-resident memory T cells, Tregs, and IFNγ- and Th17-driven transcriptional programs within the intestinal microenvironment [10, 11]. However, these models largely describe the inflammatory state once colitis is established, rather than the upstream host determinants that predispose certain patients to severe intestinal toxicity following ICI exposure.

One underexplored possibility is that pretreatment humoral immunity contributes to toxicity risk. The endogenous autoantibody (AAb) repertoire may not only reflect autoreactivity but also encode preexisting immune states that shape susceptibility or resistance to inflammation [12, 13]. This framework is particularly relevant to irC because pretreatment humoral states associated with severe toxicity may coexist with treatment-reactive immunity [14, 15]. Consistent with this possibility, we previously found that baseline serum AAb signatures were associated with both treatment outcomes and occurrence of irAE [15, 16]. In this study, baseline serum profiling identified a distinct AAb landscape associated with severe irC (sirC), characterized by reduced reactivity against immune- and mucosal-regulatory targets—including the chemokine receptors CXCR2, CXCR4, CXCR6, and CCR5—relative to patients who did not develop severe irC (nirC). These findings suggest that susceptibility to toxicity may reflect not only the emergence of proinflammatory programs, but also the absence of compensatory humoral immune states, and support the idea that baseline immunoglobulin G (IgG) AAb may help determine whether checkpoint blockade culminates in severe tissue inflammation.

A plausible mechanism by which AAb contribute to irAEs is through Fc-gamma receptor (FcγR) signaling, a central interface between humoral and innate immunity [17]. IgG immune complexes engage FcγRs on myeloid and other effector cells, triggering cytokine release, cellular activation, and tissue inflammation [18, 19]. The IgG-FcγR axis is a well-established mediator of autoimmune and gastrointestinal inflammatory disease [20, 21], yet its role in irC remains undefined. Given the mechanistic overlap between irAEs and autoimmune conditions [2, 5], FcγR-dependent immunity represents a compelling but largely untested pathway in checkpoint toxicity.

Here, we test whether humoral immunity contributes functionally to susceptibility to irC and nominate the IgG– FcγR axis as a candidate pathway of ICI-induced intestinal inflammation. Using humanized FcγR (hFcγR) mice that recapitulate IgG-FcγR interactions [22, 23], we demonstrate that transfer of polyclonal IgG from melanoma patients with sirC potentiates intestinal inflammation in the setting of PD-1 or CTLA-4 blockade, whereas wild-type mice develop minimal pathology. This response is accompanied by lymphocytic infiltration, goblet cell loss, systemic cytokine induction, and an IgG-regulated inflammatory network comprising IFNγ-producing innate-like CD8^low^ Ikzf2^high^, Itgae^high^, Itga1^high^, IL1β+ M1 macrophages, IgA/IgG plasmablast /plasma cells, and IL-22-expressing ILC3/LTi-like cells. Together, these findings position humoral immunity as a previously underappreciated determinant of irC and provide a framework for biomarker validation and mechanistic investigation in ICI-associated toxicity.

## RESULTS

### Baseline Serum AAb Profiles Define Pretreatment Humoral State Associated with irC Severity

To test whether preexisting humoral immunity marks susceptibility to irC, we established an integrated translational framework linking baseline serum AAb profiling in melanoma patients treated with ICIs to downstream mechanistic studies using patient-derived IgG and tissue-based analyses of human and experimental irC (**Fig. 1A**). Severe irC (sirC) was defined as grade 3–4 colitis, whereas grade 1–2 colitis was categorized as non-severe irC (nsirC). We used a discovery cohort from the CheckMate 915 trial (n=149) [3] to define baseline AAb profiles associated with subsequent sirC (n=31) versus nsirC (n=118). We used an independent cohort (n=18) for IgG purification and mechanistic mouse studies, to orthogonally evaluate the AAb landscape in a distinct clinical setting. In this cohort, we compared AAb profiles among sirC, nsirC, and healthy control (HC) subjects (**Fig. 1A**). **Tables S1** and **S2** summarize clinical characteristics.

**Figure 1.**
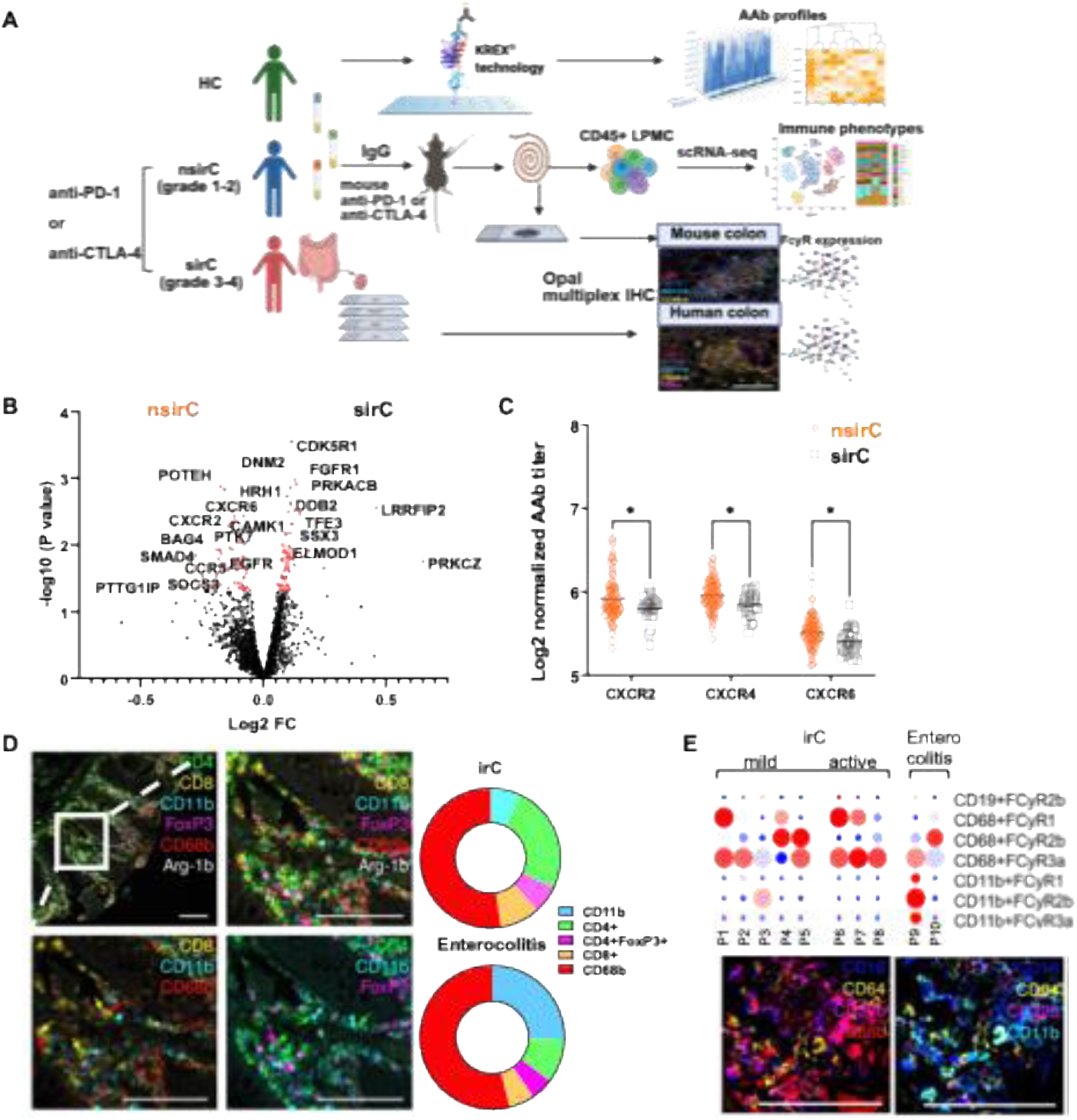
Integrating serum AAb profiling and colonic immune features in melanoma patients with *in vivo* modeling of irC. **A.** Schematic overview of the study design. We analyzed serum samples from melanoma patients treated with ICI for AAb reactivity using the KREX™ protein microarray. We transferred purified IgG from selected patients into hFcγR mice to assess how patient-derived IgG contributes to intestinal inflammation. We characterized colonic immune responses in mice using spatial proteomics and single-cell RNA sequencing (scRNA-seq) of CD45+ lamina propria mononuclear cells. We integrated these data with multiplex imaging analyses of human colonic biopsies from patients with sirC and nsirC to define immune composition and FcγR expression. Schematic created with BioRender. **B.** Volcano plot showing baseline serum AAb profiles differing between patients with sirC and nsirC. Data represent log2-normalized signal intensities; red dots indicate AAbs with significantly different titers between groups. **C.** Baseline AAb titers against selected chemokine receptors compared between patients with sirC and nsirC. Data were analyzed using multiple unpaired two-tailed *t*-tests. * p<0.05. **D.** Representative images of immune phenotypes in FFPE colon biopsies from patients with irC. Sections were stained for CD4+ (green), CD8+ (yellow), CD68A+ (red), CD11B+ (cyan), Foxp3+ (magenta), and Arg1B+ (white). Pie charts show the relative frequencies of immune cell populations in biopsies from patients with irC (*n* = 8) and enterocolitis (*n* = 2). **E.** Representative multiplex images and quantification of FcγR- expressing cells in patients with nsirC (*n* = 5), sirC (n=3), and enterocolitis (*n* = 2). The heatmap shows frequencies (%) of FcγR-expressing cell populations across colon biopsies from irC patients (P1-P10, n=10).

In the discovery cohort, global serum AAb profiles, generated using high-throughput KREX™ protein microarrays [16, 24, 25], were broadly similar across patients, arguing against irC severity being associated with generalized baseline remodeling of humoral response (**Fig. 1B**). Instead, group separation was driven by a restricted set of antigens (**Fig. 1B**; **Supplementary Fig. S1**). AAb enriched in patients with irC mapped to broadly expressed oncogenic signaling pathways, including serine/threonine kinase, MAPK, Wnt, and NF- κB/TNF-related programs: CDK5R1, FGFR1, PRKACB, LRRFIP2, DDB2, TFE3, and PRKCZ (**Table S5**). By contrast, patients protected from sirC showed relative enrichment for Ab recognizing immune-related proteins, including chemokine receptors and associated inflammatory mediators (CXCR2, CXCR4, CXCR6, CCR5, CCL5), as well as tissue-regulatory factors such as BAG4, SMAD4, CBFA2T3, and SOCS3 (**Fig. 1B**; **Supplementary Fig. S1A**). Quantitative analysis of selected candidates reinforced this pattern in nsirC relative to sirC (**Fig. 1C**). Several components of this signature have established links to intestinal inflammation *in vivo*. Disrupting CXCR2 signaling attenuates DSS-induced colitis in experimental models [27, 28], supporting a chemokine-mediated protective pathway. Similarly, SMAD4 has been implicated in constraining intestinal inflammation [26], while CBFA2T3 (MTG16) has been linked to protection from colitis via regulation of colonic epithelial differentiation, and regeneration after DSS-induced injury [27].

We next examined pretreatment AAb profiles in an independent cohort selected for functional studies. Despite clinical differences from the discovery cohort, this cohort displayed a similar central feature of baseline AAb signature: preferential enrichment of AAbs against melanoma tumor-associated and oncogenic proteins, including MLANA, MAGEA10, GAGE1, CT47A1, CDK5R1, TMEFF2, EWSR1, PRKAA and ENO1 (p<0.05; AUC>0.7) (**Supplementary Fig. S2, Table S5**). This core signature is consistent with prior reports linking tumor-associated AAbs to ICI response [15, 28, 29], an interpretation further supported by the predominance of treatment responders in this cohort (**Table S2**).

Conversely, AAb recognizing immune- or tissue-regulatory proteins were relatively depleted in patients with sirC. Anti-BRD2 titers were 6-fold lower than in HC (p< 0.001; **Supplementary Fig. S2A**), consistent with attenuation of a broader immune-regulatory humoral state given the role of BET family proteins in inflammatory transcription and T-cell differentiation [30]. Among anti-PD-1-treated patients, titers of AAb targeting HRH2 and OVOL1 were similarly reduced (**Supplementary Fig. S2A**), suggesting altered mucosal immune regulation and epithelial homeostasis [31, 32]. Anti-BASP1 AAb titers were also lower in all sirC than in nsirC (2-fold; p<0.05) and HC (12.3-fold; p < 0.001; **Supplementary Fig. S2A**), although because BASP1 has been linked to MYC-associated oncogenic signaling [33], this reduction may reflect altered tumor-associated humoral recognition in sirC rather than a regulatory axis.

Treatment-specific differences were also evident. In anti-PD-1-treated patients, sirC was associated with increased reactivity against metabolic and stress-response proteins, including AMPK, PDK4, SEPTIN4, STK17B, PPP2CB, and PRKAA2 (fold change >1.5 vs. nsirC, p<0.05, AUC>0.7; **Supplementary Fig. S2A**). In anti-CTLA-4-treated patients, sirC was instead associated with antibodies targeting apoptotic and differentiation-related proteins, including FADD, MX1, MALL, and PRKD3 (fold change >1.5 vs. nsirC, p<0.05, AUC>0.7). Across treatment groups, nsirC samples more closely resembled HC than sirC.

Together, these findings identify a selective pretreatment AAb profile in irC severity rather than nonspecific baseline autoimmunity and provided the rationale for *in vivo* testing whether patient-derived IgG can modulate colonic inflammation. Across cohorts, sirC was linked to a preserved AAb signature against tumor-associated antigens together with relative depletion of antibodies directed against immune- and tissue-regulatory proteins. These features are most appropriately interpreted as markers of a broader host immune state predisposed to intestinal toxicity, not as direct evidence for antigen-specific causality. and they provided the rationale to test directly whether patient-derived IgG can modulate colonic inflammation.

### Human irC colon biopsies reveal FcγR-expressing myeloid-rich inflammation

We next characterized inflamed colon biopsies from melanoma patients with irC to define the tissue immune context of clinical disease and inform subsequent mechanistic studies in hFcγR mice (**Fig. 1A, D, E**). The biopsy cohort included patients with mild (n=5), chronic (n=3), and entero-colitis (n=2), representing a spectrum of irC pathology (**Supplementary Fig. S3A**), but not normal or ICI-treated non-colitis mucosa.

Histopathologic examination identified canonical features of irC, including lymphoid aggregates, epithelial mucin depletion, focal cryptitis, and crypt architectural distortion [8, 34]. As a complementary systemic measure of inflammation, circulating IL-17A levels were elevated in anti-PD-1 treated patients with irC compared with HC (**Supplementary Fig. S3B**), consistent with prior work implicating inflammatory cytokine programs in checkpoint-associated intestinal toxicity [8, 35].

Multiplex immunohistochemistry (IHC) (**Supplementary Table S3**) demonstrated prominent CD68+ macrophage-rich infiltrates across biopsies, comprising approximately 40%–70% of total cells. CD4+ and CD8+ T-cells were proportionally more abundant in biopsies with less severe inflammatory pathology, comprising approximately 22% and 8% of total cells, respectively (**Fig. 1D**; **Supplementary Fig. S3C, D**). This pattern is consistent with prior single-cell analyses showing expansion of both inflammatory myeloid cells and cytotoxic proliferating T-cells during early or less severe stages of irC [10].

We next examined FcγR expression within the inflamed colon in this irC cohort to identify potential cellular targets of IgG-mediated immune activation. Expression of the activating FcγRs: CD64/FcγRI, CD32A/FcγRIIA, and CD16/FcγRIII, as well as the inhibitory FcγR CD32B/FcγRIIB (**Supplementary Table S4**), was evaluated in the colon tissue. CD16 localized predominantly on CD68+ macrophages, whereas CD64 and CD32B were detected on both CD68+ and CD11b+ myeloid populations (**Fig. 1E**; **Supplementary Fig. S3E**). In contrast, CD32B+CD19+ B cells were confined to lymphoid aggregates and detected in only 2 of 10 biopsies (**Supplementary Fig. S3E**). Within this irC cohort, myeloid cells expressing activating FcγR were more abundant than cells expressing inhibitory FcγR across biopsies (**Fig. 1E**; **Supplementary Fig. S3E**), consistent with a predominance of activating FcγR signaling within inflamed mucosa [19].

Together, these findings identify intestinal myeloid cells as the principal FcγR-expressing population in irC and demonstrate the prevalence of activating FcγR expression within inflamed tissue. These data support a model in which early or less tissue-destructive irC represents a niche enriched for T cells and FcγR-competent myeloid cells that can respond to IgG-mediated immune signals, providing a rationale for testing the contribution of patient-derived IgG in hFcγR mice.

### Patient-derived IgG Amplifies Fcγ Receptor-Dependent Colon Inflammation in ICI-treated mice

To test whether patient-derived IgG contributes to ICI-associated intestinal inflammation [36], we transferred purified serum IgG from melanoma patients with sirC, nsirC, or HC into C57BL/6-hFcγR mice (2–6 mice per IgG donor) [22] and, in selected experiments, WT C57BL/6 mice used as controls (**Fig. 2A**). Mice subsequently received anti-PD-1 or anti-CTLA-4 monotherapy matched to the ICI class received by the IgG donor, or the corresponding isotype control. Additional control groups included hFcγR mice receiving IgG alone without checkpoint blockade and untreated hFcγR mice. We monitored mice longitudinally for clinical signs of toxicity and evaluated them at study completion for intestinal, hepatic, and systemic inflammatory responses. C57BL/6 control mice, which received the same IgG and ICI treatments as hFcγR mice, exhibited no significant histopathologic evidence of intestinal inflammation (**Supplementary Fig. S4B, D**). In contrast, hFcγR mice administered sirC IgG in combination with anti-PD-1 or anti-CTLA-4 developed mild intestinal inflammation, characterized by increased lamina propria (LP) cellularity, mucosal hyperplasia, and prominent submucosal leukocytic infiltration [5, 7] (**Fig. 2B-D**; **Supplementary Fig. S4C, D**). Although the inflammatory phenotype was relatively mild, the lesions reproduced key histopathologic features reported in both patients with irC and established murine irC models [10, 39]. Accordingly, histologic colitis scores [38] were significantly higher in sirC IgG mouse group than in the nsirC IgG mouse group under anti-PD-1 treatment (p=0.015, Mann-Whitney) (**Fig. 2C**). In anti-CTLA-4 treatment arm, sirC IgG transfer resulted in higher leukocyte infiltration scores than HC IgG, though this difference did not reach statistical significance (p=0.076), consistent with the limited number of independent HC donors available in this arm (n=3); the same direction of effect was observed for sirC versus nsirC IgG under anti-CTLA-4 treatment (p=0.79). Consistent with the limited severity of intestinal inflammation, hFcγR mice showed no evidence of acute systemic toxicity, including alopecia, gastrointestinal bleeding, or significant weight loss (**Supplementary Fig. S4A**).

**Figure 2.**
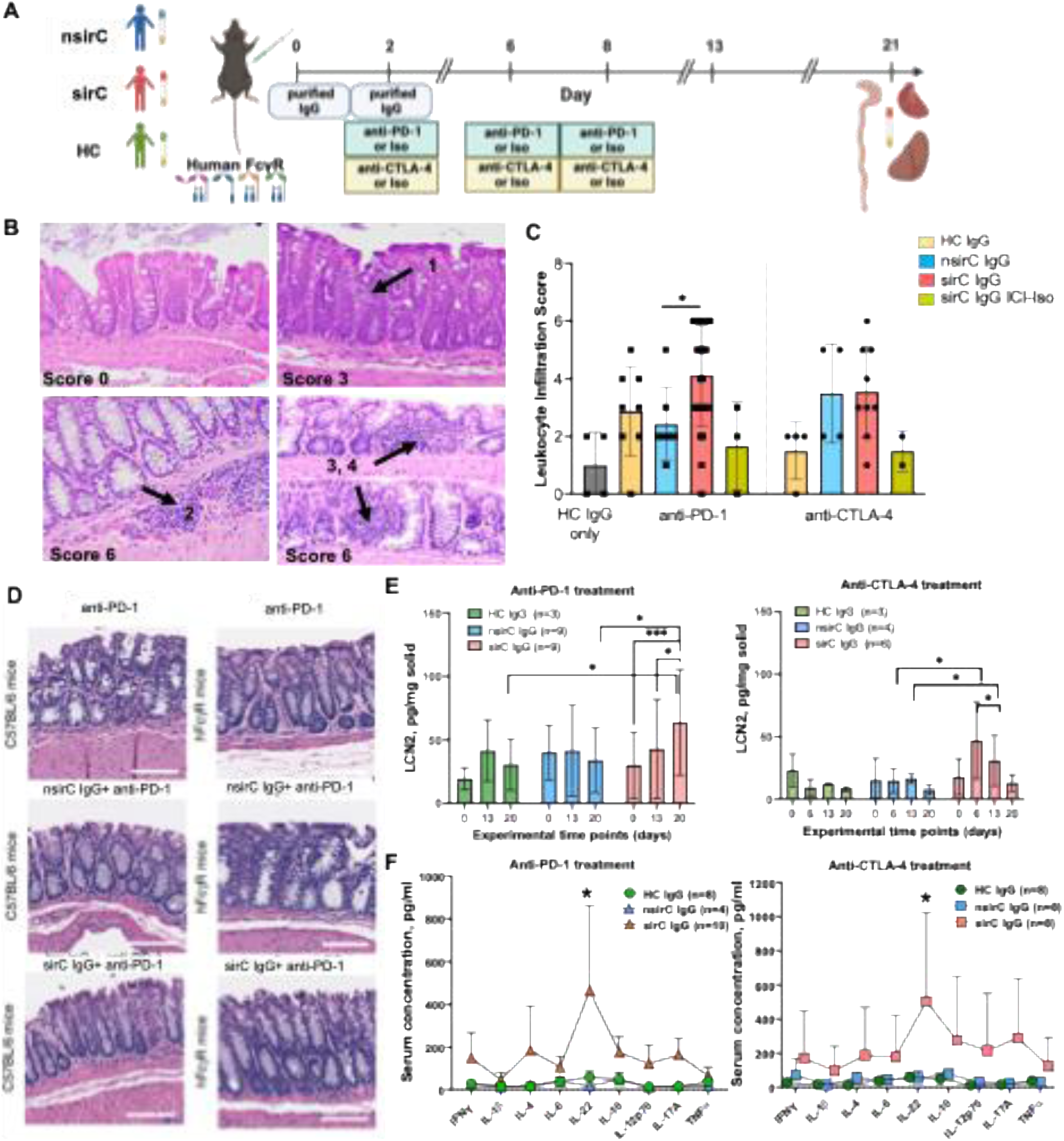
*In vivo* modeling of irC using patient-derived IgG in FcγR-humanized mice. **A.** Experimental design: IgG was purified from serum of melanoma patients with nsirC, sirC, or HC and administered intraperitoneally into hFcγR mice (≥2 mice per patient). Mice received anti-PD-1 or anti-CTLA-4 antibodies (200 µg on days 2, 6, and 8) or isotype controls. Mice were monitored for weight loss and clinical and euthanized on day 21 for blood and tissue (colon, liver, and heart) collection. **B.** Representative H&E-stained colon sections (40X magnification) with corresponding histologic scores for leukocyte infiltration (scale 0–6), illustrating minimal (score 0) versus severe (score 6) infiltration. **C.** Quantification of colonic leukocyte infiltration across experimental groups (each dot represents an individual mouse; 2–6 mice per IgG donor), assessed in a blinded fashion. A donor-level analysis is shown in **Supplementary Fig. S5A**. **D.** Representative H&E-stained colon sections (scale bar 100 µm) from C57BL/6 and hFcγR mice receiving anti-PD-1 alone and together with nsirC or sirC IgG, illustrating greater colonic pro-inflammatory changes in hFcγR mice with no substantial changes in C57BL/6 controls. **E.** Fecal lipocalin-2 (LCN2) concentrations over time in anti-PD-1- and anti-CTLA-4-treated mice. **F.** Differential serum cytokine levels in sirC versus nsirC IgG recipients under anti-PD-1 (sirC n=9, nsirC n=4) and anti-CTLA-4 (sirC n=8, nsirC n=8) treatment. Each point represents one of nine cytokines (IFN-γ, IL-4, IL-6, IL-1β, IL-17A, IL-22, IL-12p70, IL-10, TNF-α) measured by a multiplex bead assay. Statistical analysis: **C.** Two-tailed Mann-Whitney test. **E.** Two-way ANOVA with Tukey’s post-hoc test comparing time points within each IgG group. **F.** Two-tailed Mann-Whitney test with Benjamini-Hochberg correction for multiple comparisons across the 9-cytokine panel. Error bars represent mean ± SD. *p < 0.05, **p < 0.01

Because histologic scoring included multiple recipient mice per IgG donor, we additionally examined this comparison at the donor level, averaging histology scores across all recipient mice for each donor (**Supplementary Fig. 5A**). At the donor level, sirC IgG donors showed a consistently higher mean histology score than nsirC or HC IgG donors across both the anti-PD-1 and anti-CTLA-4 arms, with the same direction of effect observed in every comparison as in the mouse-level analysis. Given the reduced number of independent donors available for this analysis (3–11 donors per group; **Supplementary Table S2**), this donor-level comparison serves as a robustness check of the primary mouse-level result rather than an independently powered analysis. This limitation was greatest in the anti-CTLA-4 arm, which included only three donors per group; those findings should therefore be considered preliminary.

In hFcγR mice, sirC IgG treatment was associated with transient increases in fecal lipocalin-2 (LCN2), a marker of intestinal inflammation, under both ICI regimens, although levels remained below those typically associated with acute colitis (100 pg/mg) [40] (**Fig. 2E**). Peak LCN2 occurred on day 6 following anti-CTLA-4 and on day 20 following anti-PD-1 treatment, supporting a mild intestinal inflammatory phenotype. In the anti-PD-1 cohort, sirC IgG treatment was also associated with leukocyte infiltration in the liver and increased serum alanine aminotransferase (ALT) levels [41], consistent with mild hepatic inflammatory injury; these findings were not observed with anti-CTLA-4 plus sirC IgG (**Supplementary Fig. S5B**).

Serum cytokine profiling further demonstrated a broad inflammatory signature in sirC IgG recipient mice (**Fig. 2F**). Under anti-PD-1 treatment, sirC IgG recipients showed significantly higher levels of IL-17A and IL-10 relative to nsirC IgG or HC IgG (q=0.003 and q=0.005, Mann-Whitney test with Benjamini-Hochberg correction for multiple comparisons) (**Supplementary Fig. S5C**). IL-22 showed the largest relative increase, with mean serum concentrations approximately 10-fold (q<0.05) (**Fig. 2F**). Under anti-CTLA-4 treatment, cytokine levels in sirC IgG recipients showed the same directional trend relative to nsirC IgG recipients, mostly for IL-22 (mean fold-change approximately 9-fold), but did not retain significance after correction for multiple comparisons (**Fig. 2F**), consistent with the smaller and more variable anti-CTLA-4 cohort noted elsewhere in this study. The observed cytokine profile was consistent with serum IL-17 levels in melanoma patients with irC, as reported previously [2, 42–44]. In contrast, IL-22 was not detectable in serum from melanoma patients with irC, suggesting that IL-22 may be more relevant to an early phase of inflammation in this model than to established human irC. The Th22-related cytokine signature observed in hFcγR mice is otherwise consistent with prior studies in intestinal inflammation [45–47].

We next examined whether changes in the intestinal microbiota accompanied IgG- and ICI-associated inflammation. Longitudinal 16S rDNA sequencing did not reveal significant differences in overall microbial composition during the 21-day experiment (**Supplementary Fig. S5D**). These findings indicate that major microbiome remodeling was not required for the early inflammatory phenotype observed in this model.

Together, these findings indicate that patient-derived IgG functions as a disease-modifying factor that amplifies susceptibility to ICI-associated intestinal inflammation rather than acting as an autonomous driver of inflammation. The hFcγR mouse model provides an experimental framework to define the cellular and molecular mechanisms linking baseline humoral immunity to irC.

### Patient-Derived IgG and Checkpoint Blockade Shape Distinct Colonic Immune Programs in Humanized **FcγR Mice**

To test whether hFcγR mice recapitulate key immune features of irC, we examined colonic pathology and immune phenotypes after transferring patient-derived IgG in combination with anti-PD-1 or anti-CTLA-4 treatment. Histology revealed dense leukocytic infiltrates reminiscent of tertiary lymphoid structures, although the staining panel was insufficient to establish this identity definitively (**Fig. 3A**; **Supplementary Fig. S6A-C**). Multiplex imaging showed spatial organization of these inflammatory foci with readily detectable CD4+ and CD8+ T cells, CD11b+ myeloid cells, F4/80+ macrophages, and CD4+Foxp3+ Treg cells (**Fig. 3A**). Image-based quantification suggested that nsirC IgG lesions were relatively enriched for CD11b+ and F4/80+ myeloid cells, whereas sirC IgG leukocyte lesions showed greater T-cell (CD4+ and CD8+) representation; anti-PD-1 treated colons favored F4/80+ macrophages, whereas anti-CTLA-4 treated colons showed relatively greater CD11b+ myeloid abundance. Treg cells localized preferentially near T cells and F4/80+ macrophages, supporting distinct inflammatory architectures in checkpoint contexts (**Supplementary Fig. S6D**).

**Figure 3.**
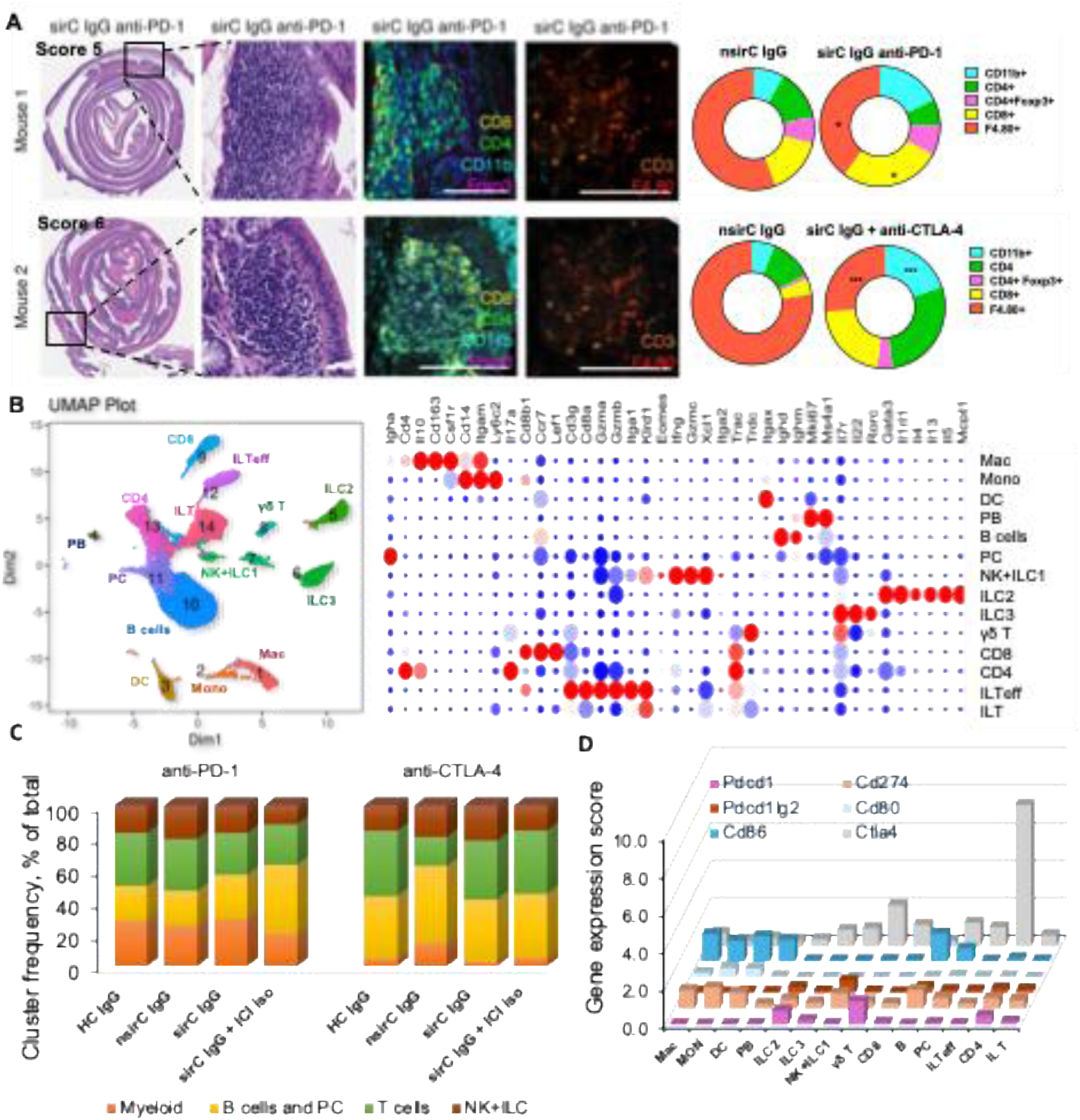
Humanized FcγR mice recapitulate colonic immune phenotypes associated with irC. **A.** Representative histologic and immune phenotypic features in FFPE colon sections from hFcγR mice following modeling of immune-related colitis with immune checkpoint blockade and patient-derived IgG transfer. H&E images show representative colonic pathology and corresponding inflammation scores. We performed multiplex immunofluorescence staining using the same marker panel as in human colon biopsies, substituting F4/80 for CD68 to account for species-specific differences in macrophage detection. Colon sections were stained for CD8 (yellow), CD4 (green), F4/80 (red), CD11b (cyan), and Foxp3 (magenta). Donut plots summarize immune cell frequencies (%) in the mouse colon across experimental groups. **B.** UMAP visualization of major CD45+ immune cell populations identified in the colonic LP across all samples. The accompanying bubble plot shows expression of key marker genes used for cluster annotation, with dot size representing the proportion of cells expressing each gene and color indicating average expression level. **C.** Relative abundance of major immune cell populations in the colonic LP of hFcγR mice treated with anti–PD-1 or anti–CTLA-4 and receiving patient-derived IgG. Stacked bar plots show the proportional representation of each immune subset across treatment groups. **D.** Distribution of immune checkpoint receptors and ligands across colonic immune cell subsets identified by scRNA-seq.

To define the underlying immune programs, we performed scRNA-seq on CD45+ LP cells from hFcγR mice following transfer of patient-derived IgG and treatment with anti-PD-1 or anti-CTLA-4. Because we pooled samples by subgroup, these data reflect condition-level immune states rather than inter-mouse variability; we also performed anti-PD-1 and anti-CTLA-4 experiments separately, so we focused on within-treatment comparisons. Across pooled subgroup samples, 73,078 CD45+ cells resolved into 14 immune populations spanning myeloid, innate lymphoid, T-cell, and B-cell compartments (**Fig. 3B, C**). These included macrophages (Mac; *Cd14, Cd163*), dendritic cells (DC; *Cd11c*), monocytes (Mon; *Ly6c*), innate lymphoid (ILC) populations including ILC1/NK-like cells (*Xcl1, Klrd1, Ifng, Prf1*), ILC2 (*Gata3, Il4, Il5, Il13*), and ILC3 (*Rorc, Il22*). We identified multiple B-cell states (*Cd19*): naïve B cells (*Ccr7*), germinal center (GC-like, *Mki67*), and plasma cells (PC) expressing immunoglobulin transcripts. Distinct T-cell populations were also identified, including naïve CD4 and CD8 T cells (*Ccr7*), γδ T cells (*Trdc*), and innate-like T (ILT; *Cd8a*^low^, *CD160, Kit)*, including the effector (ILTeff) subset (*Gzmb, Gzmc, Itga1^high^*) (**Fig. 3B**; **Supplementary Fig. S7; Table S5**). In contrast to microbiota-driven irC models [10, 39] in which neutrophilic inflammation is prominent, neutrophils (*Cxcr2*) accounted for only a minor fraction of CD45+ LP cells (0.05%) in our model and were excluded from downstream analyses. This difference may reflect the modest severity and early stage of inflammation observed following patient-derived IgG transfer, or it may suggest that FcγR-dependent mechanisms preferentially shape lymphoid and mononuclear cell responses.

Overall, checkpoint context was a dominant determinant of the colonic immune landscape (**Fig. 3C**; **Supplementary Figs. S8A, 8B**). Anti-PD-1-treated colons were relatively enriched for myeloid populations (>20% of CD45+ cells), whereas anti-CTLA-4-treated colons showed greater representation of innate lymphoid populations, particularly ILC (∼10% of CD45+ cells), as well as ILT populations (∼5% of CD45+ cells). (**Fig. 3C**; **Supplementary Fig. S8A, S8B**). Principal component analysis further supported segregation by treatment context (**Supplementary Fig. S8C**). the Mac patient-derived IgG did not induce a single stereotyped inflammatory state, but instead amplified checkpoint-dependent immune programs in the colon. Consistent with this interpretation, checkpoint-related transcripts were differentially distributed across immune compartments (**Fig. 3D**). *Pdcd1* expression was enriched in ILC2, γδ T cells, and selected CD4+ T-cell populations, whereas *Ctla4* expression was most prominent in plasma cells and was also detected in γδ and CD4+ T-cell populations. In contrast, transcripts encoding checkpoint ligands and costimulatory molecules, including *Cd274* (*Pd-l1*), *Pdcd1lg2* (*Pd-l2*), *Cd80*, and *Cd86*, were concentrated in myeloid and B-cell compartments. Other inflammatory mediators showed broadly similar cellular distributions across treatments (**Fig. 3B**), including *Gzma* and *Gzmb* in ILTeff, *Gzmc* in ILC1/NK-like cells, *Ifng* in ILC1 and selected CD4+ T- cell subsets, and *Il6* primarily in monocytes. A shared feature across both ICI settings was increased *Il22* expression within ILC3 cells, suggesting convergence on a common mucosal inflammatory program despite differences in the dominant immune compartments.

Together, these data establish the hFcγR model as a tractable system for studying IgG-dependent mechanisms of irC and identify checkpoint-specific immune architectures that motivated higher-resolution analysis of lymphocyte states.

### T- and B-cell Programs Associated with ICI- and IgG-dependent Colonic Inflammation in hFcγR mice

Because PD-1 and CTLA-4 primarily regulate lymphocyte activation, we next profiled T- and B-cell compartments at higher resolution. Re-clustering of CD3+ cells identified 10 transcriptionally distinct T-cell states in the colonic LP (**Fig. 4A, B**). Overall, T-cell composition was broadly preserved across treatment groups, with only modest shifts in relative abundance (**Fig. 4C, D**). Most cells (40%–60%) were distributed across naïve CD4/CD8 (*Ccr7, Sell*) and ILT (*Cd103, Cd8a, Cd8b, Cd160*) states, whereas the remaining clusters represented activated effector and regulatory populations [48].

**Figure 4.**
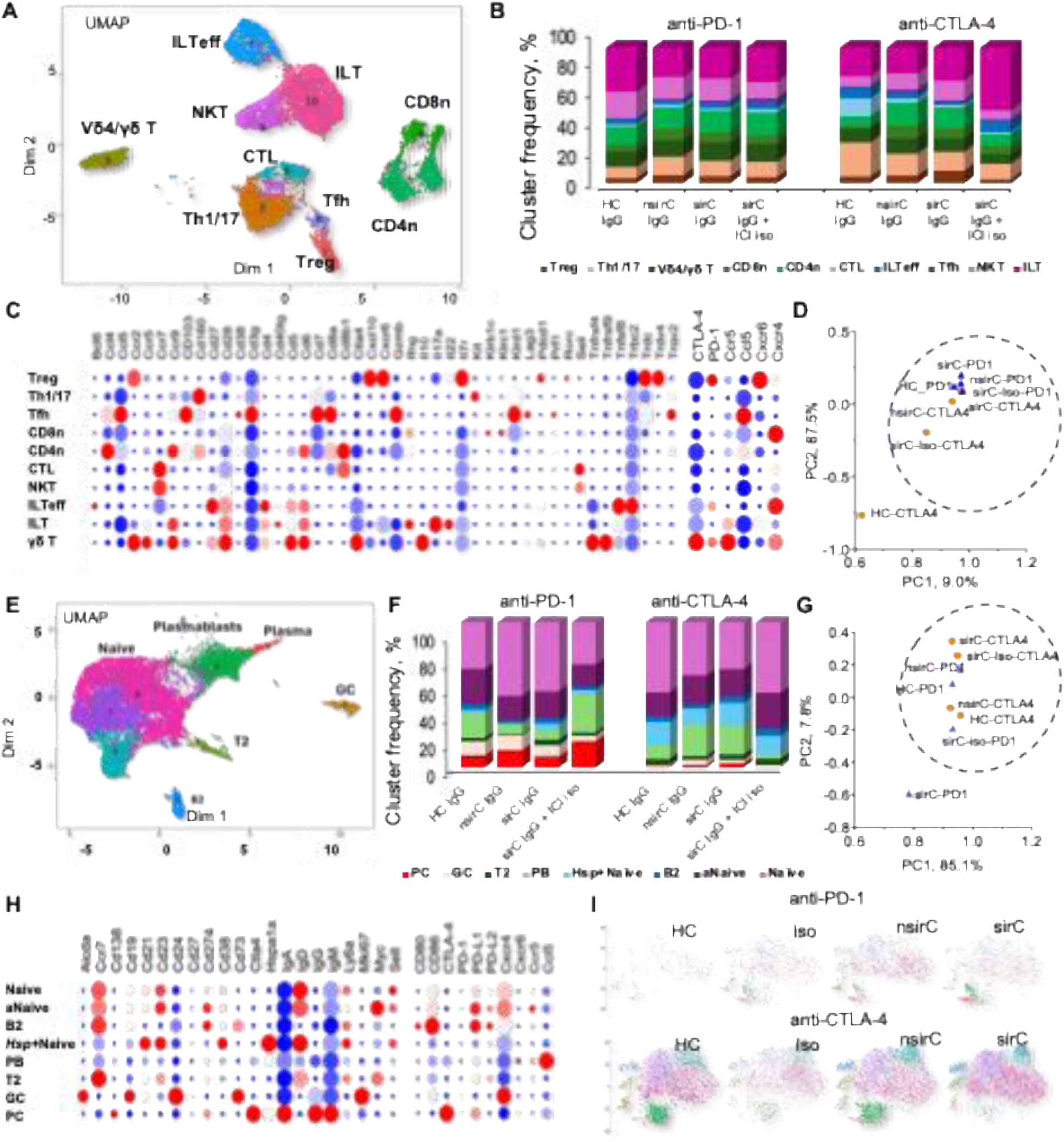
Ab- and checkpoint-mediated remodeling of T and B cell compartments in the colonic LP of hFcγR mice. **A.** UMAP visualization of 10 transcriptionally distinct CD3+ T cell clusters identified in the colonic LP across all samples. **B.** Relative frequencies of CD3+ T cell clusters across treatment groups, shown as proportional representation (%) of each cluster within the colonic LP. **C.** Bubble plot showing expression of key marker genes used for annotation of CD3+ T cell clusters. Dot size represents the proportion of cells expressing each gene, and color indicates average expression level. **D.** PCA based on the relative frequencies of CD3+ T cell clusters across treatment groups. The PC1 versus PC2 plot shows the sample distribution and grouping based on differences in T cell composition. **E.** UMAP visualization of 8 transcriptionally distinct CD19+ B cell clusters identified in the colonic LP across all samples. **F.** Relative frequencies of CD19+ B cell clusters across treatment groups, shown as proportional representation of each cluster within the colonic LP. **G.** PCA based on the relative frequencies of CD19+ B cell clusters across treatment groups. The PC1 versus PC2 plot shows the sample distribution and grouping based on differences in B cell composition. **H.** Bubble plot showing expression of key marker genes used for manual annotation of CD19+ B cell clusters. Dot size represents the proportion of cells expressing each gene, and color indicates average expression level. **I.** Individual t-SNE maps showing the distribution of CD19+ B cell populations for each treatment group.

Among activated populations, Th1/Th17-like (*Ccr9, Cd40lg*), NKT-like (*Klrc1*), and cytotoxic T-cell (CTL) (*Cd8a, Cd8b, Ccl4, Ccr9*) clusters expressed inflammatory programs, including *Ifng* and/or *Il17a*, indicating multiple potential sources of type 1 inflammatory signaling in the colon (**Fig. 4B, E**). We also identified an activated Treg population marked by *Il10, Tnfrsf4*, *Tnfrsf9*, *Ccr2*, *Cxcr4*, *Ccr9*, and *Ctla4*, and costimulatory receptor expression, with comparatively lower *Pdcd1* than other subsets (**Fig. 4B, E**); this population trended higher in anti-CTLA-4 treated mice receiving sirC IgG (**Fig. 4C**). Tfh (*Cd27, Bcl6, Tnfsf8, Pdcd1*) and γδ T-cell (*Trdc, Trgc, Rorc, Cxcl10, Cxcr6, Il7r*) clusters co-expressed *Pdcd1* and *Ctla4*, whereas Th1/Th17-like cells showed a *Ctla4*-skewed profile (**Fig. 4E**). *Ccl5* expression was most prominent in ILTeff, whereas *Cxcr4* expression was detected across activated T-cell clusters, except Th1/Th17-like cells and ILT (**Fig. 4H**).

We next analyzed CD19+ cells, given the prominence of B-cell populations in the global dataset and the organized aggregates seen histologically. Re-clustering identified 8 transcriptionally distinct B-cell states, including follicular naïve (*Ighd, Ighm, Ccr7, Sell, Fcer2a*), transitional T2-like (*Sell^low^*), GC-like (*Nt5e, Aicda, Mki67*), plasmablast, and PC populations (**Fig. 4E-G**). As in the T-cell compartment, overall B-cell composition was largely similar across groups (**Fig. 4F, G**). The clearest difference involved antibody-secreting populations: under anti–PD-1 treatment, mice receiving sirC IgG showed expansion of plasmablasts and PC relative to the other IgG groups, whereas these populations were markedly reduced under anti–CTLA-4 treatment, irrespective of IgG source (**Fig. 4F, G**).

Checkpoint-related gene expression further distinguished B-cell states (**Fig. 4H**). Checkpoint ligands and costimulatory molecules were most evident in naïve and B2-like populations, whereas plasmablasts and PC expressed *Ctla4* with minimal *Pdcd1*. Notably, these antibody-secreting cells were the only B-cell populations with appreciable *Ctla4* expression, raising the possibility that they are selectively perturbed under anti-CTLA-4 conditions and contribute to the distinct colonic immune architecture induced by CTLA-4 blockade.

Overall, high-resolution lymphocyte profiling showed that patient-derived IgG and checkpoint blockade remodel discrete effector, regulatory, and antibody-secreting states rather than global lymphocyte repatterning. These data define candidate cellular circuits associated with sirC IgG-driven intestinal inflammation and further support the conclusion that anti-PD-1 and anti-CTLA-4 elicit partially overlapping but mechanistically distinct mucosal immune programs.

### FcγR-expressing Myeloid Cells Translate sirC IgG into Checkpoint-specific Inflammatory Programs

Given the prominence of FcγR-expressing myeloid cells in human irC biopsies, we next asked if this compartment in hFcγR mice was positioned to integrate patient-derived IgG with checkpoint-driven inflammatory cues. Across treatment groups, inflammatory cytokine expression mapped to a limited set of cellular compartments, with myeloid cells, CD4 T cells, and ILC populations comprising the dominant cytokine-producing subsets in the LP (**Fig. 5A**). *Il1b* localized primarily to monocyte/macrophage populations, *Il17*a to CD4 T-cell subsets, and *Il22* predominantly to ILC3 cells. Increased serum Il22 levels (**Fig. 2F**) emerged as a shared feature of sirC IgG-associated inflammation across both checkpoint settings, identifying the IL-22 axis as a shared correlate of early mucosal remodeling despite otherwise distinct immune architecture, consistent with activation of mucosal barrier-associated inflammatory pathways that have been implicated in experimental and human intestinal inflammation [10, 39].

**Figure 5.**
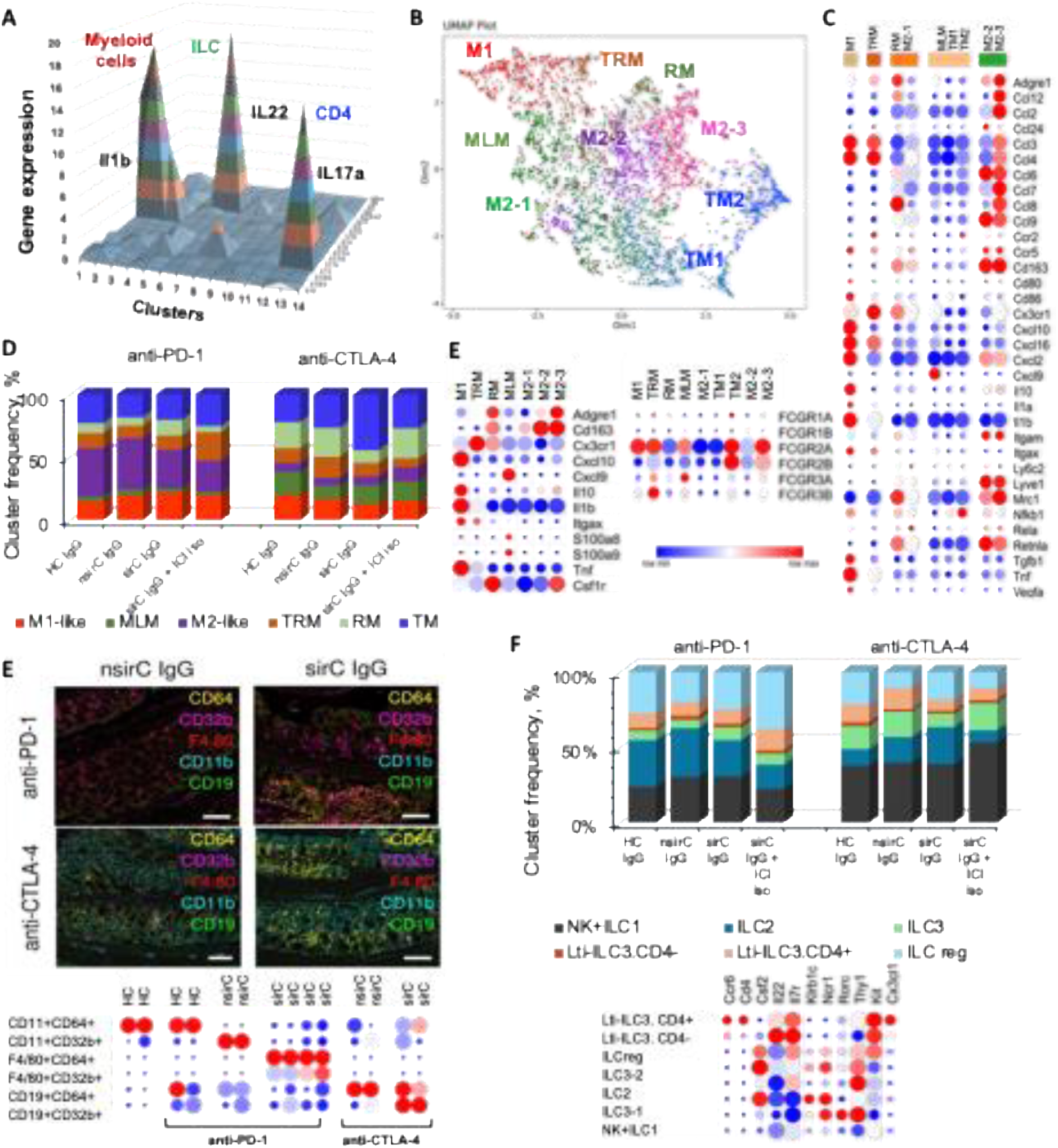
Myeloid and ILC remodeling in the colonic lamina propria identifies FcγR-expressing macrophage states and IL-22-associated innate programs in hFcγR mice. **A.** Cell-type-resolved cytokine expression landscape of the colonic lamina propria, showing the major cytokine- producing immune populations and their relative transcriptional contributions. Myeloid cells, ILC3/LTi-like cells, and CD4 T cells comprise the dominant inflammatory cytokine-expressing compartments. **B.** UMAP projection of colonic macrophages showing transcriptionally distinct subclusters identified by unsupervised clustering, with annotation of inflammatory (M1-like), tissue-resident (TRM), monocyte-like (MLM), M2-like states and remodeling/transitional populations (RM and TM). The bubble plot shows expression of representative marker genes across macrophage subclusters. Dot size indicates the fraction of cells expressing each gene, and color indicates average expression. **C.** Relative frequencies of macrophage subclusters across treatment groups, shown as the proportional representation of each subset within the Mac compartment. **D.** Expression of key immune-regulatory genes and human Fcγ receptors across colonic macrophage subclusters. **E.** Representative multiplex immunofluorescence images of colon sections from hFcγR mice showing CD64, CD32b, F4/80, CD11b, and CD19 under anti-PD-1 or anti-CTLA-4 treatment following transfer of patient-derived IgG, with quantification of FcγR-expressing myeloid populations across treatment groups. **F.** Relative frequencies of ILC3 subclusters across treatment groups, shown as the proportional representation of each subset within the total ILC compartment. **G.** Bubble plot shows expression of representative marker genes used to annotate major ILC populations. Dot size indicates the fraction of cells expressing each gene, and color indicates average expression.

To identify candidate IgG-responsive effector populations, we subclustered colonic *Adgre1+* macrophages and identified nine transcriptionally distinct states, including inflammatory, resident-like, monocyte-derived, and remodeling/transitional (RM and TM) populations (**Fig. 5B, D**). Inflammatory subsets, including M1-like (*Il1b, Tnf, Cxcl10, Itgax, Cd86*), tissue-resident (TRM; *Cx3cr1, C1qa*), and monocyte-like (MLM) (Il1b, Tnf, Cxcl9/Cxcl10, S100a8, S100a9), were enriched for activating human FcγRs, particularly *Fcgr2a*, with additional *Fcgr3* expression in TRM and MLM populations. In contrast, expression of the inhibitory receptor *Fcgr2b* was relatively preserved in M2-like (*Cd163, Il10*) populations, which differed in activation marker expression (*Cd163, Mrc1, Retnla, Ccl6, Ccl8*) (**Fig. 5B, D**). Transcriptional profiling of mouse colonic macrophages revealed upregulation of complement genes (*C1qa, C1qb, C1qc*), consistent with their dual roles in immune complex clearance and inflammation amplification [23, 49]. Thus, the Mac subsets most enriched for inflammatory mediators also exhibited the most activating FcγR configuration, identifying them as likely responders to pathogenic IgG.

The distribution of Mac subpopulations in mouse colon differed by checkpoint context. Anti-PD-1-treated colons showed greater overall myeloid representation than anti-CTLA-4-treated colons. In this setting, sirC IgG preferentially expanded inflammatory M1-like, RM, and TRM subsets (**Fig. 5C, E**), which single-cell multiomics has recently linked to human and *in vivo* models of irC [10, 39]. In contrast, anti-CTLA-4-treated colons contained fewer Mac overall and had a lower frequency of M2-like populations than anti-PD-1-treated colons; instead, MLM-like and TM populations were more abundant and increased further after transfer of sirC IgG, suggesting a distinct inflammatory architecture despite partially overlapping cytokine outputs. Multiplex immunofluorescence supported these findings at the tissue level: CD64 and CD32b localized predominantly to F4/80+ macrophages and CD11b+ myeloid cells, with FcγR-expressing F4/80+ macrophages most evident in anti-PD-1 treated mice receiving sirC IgG, whereas under anti–CTLA-4 FcγR signal was observed primarily in CD11b+ myeloid cells (**Fig. 5E**). These data indicate that PD-1 and CTLA-4 blockade engage distinct FcγR+ myeloid populations in the colon.

Because Il22 was a shared cytokine feature across treatment groups (**Fig. 2F**), we next resolved the colonic ILC compartment. Sub-clustering identified transcriptionally distinct ILC populations based on canonical markers (**Fig. 3B**) and subcluster-specific, including ILC1/NK-like cells expressing *Xcl1, Klrd1, and Ifng*; cytotoxic NK cells additionally expressing *Prf1*; ILC2 cells expressing *Gata3, Il4, Il5, and Il13*; ILC3 cells expressing *Rorc* and *Il22*; and an LTi-like ILC3 population expressing *Ccr6*, *Cx3cl1, Il22,* and *Il23r* (**Fig. 5F**). Although the overall ILC3 compartment was preserved across groups, subset-level shifts were evident in both proportional abundance within the ILC3 pool and contribution to total CD45+ cells (supplementary **Fig. S9D**). These data place ILC3s as a major source of mucosal IL-22 in this model and suggest that, in contrast to the more divergent remodeling of FcγR+ myeloid populations under different ICIs, IL-22-producing innate lymphoid programs may represent a more convergent axis of sirC IgG-associated inflammation.

To define how patient-derived IgG reshaped the inflammatory milieu within each checkpoint context, we compared LP transcriptional states in mice receiving sirC versus nsirC IgG under anti-PD-1 or anti-CTLA-4 treatment. sirC IgG induced distinct inflammatory programs in the two ICI settings (**Fig. 6A**): under anti-PD-1, an innate inflammatory profile marked by *Il1b, Il6, Il15, Ccl3, Ccl4*, and *Tnf*; under anti-CTLA-4, a lymphoid-skewed inflammatory profile marked by *Ifngr1, Il21r, Tnf*, and *Ccl5*.

**Figure 6.**
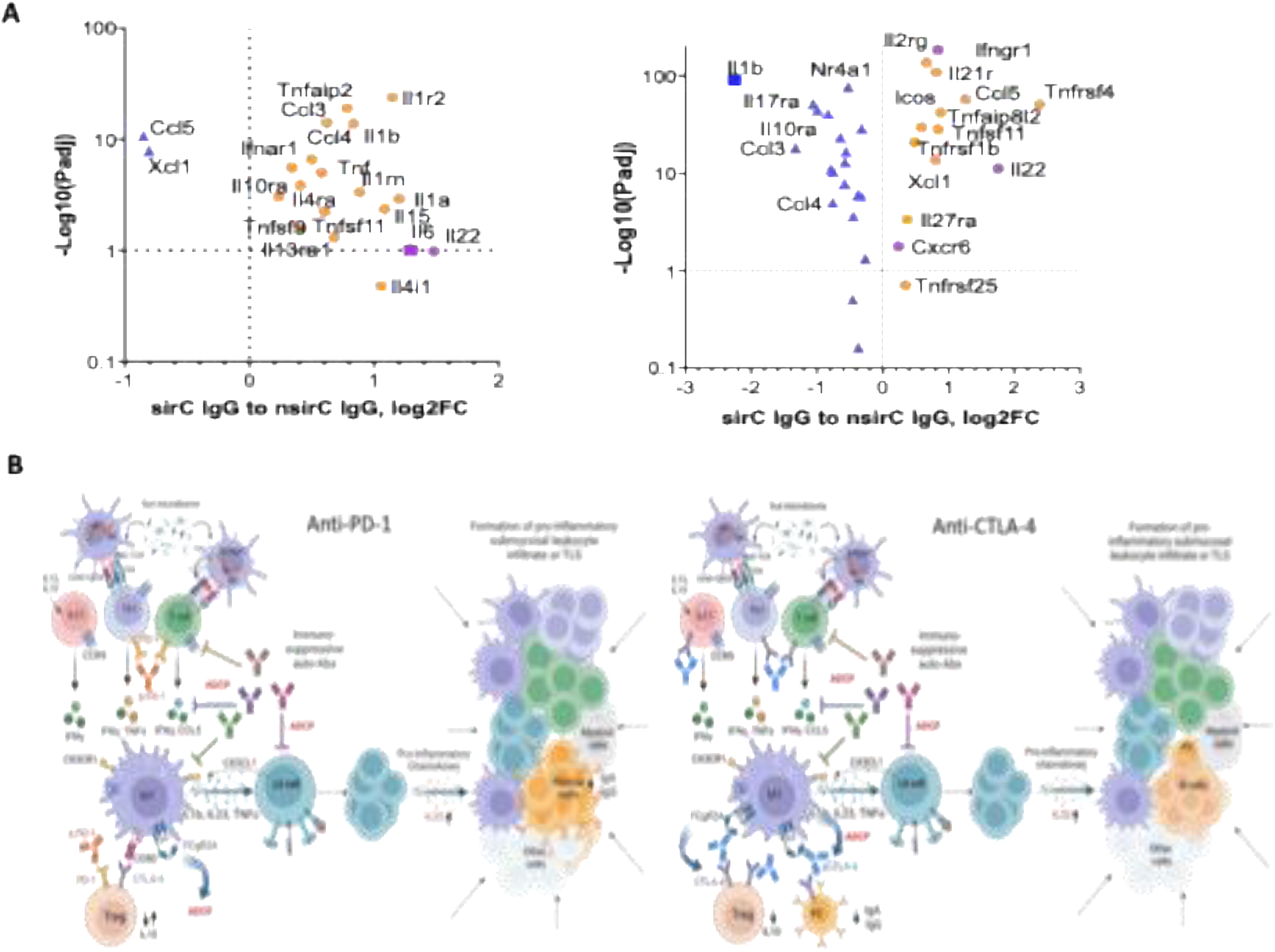
Transcriptomic remodeling of the colonic LP and proposed mechanisms underlying irC during anti-PD-1 or anti-CTLA-4 therapy. **A.** Volcano plots showing differential gene expression in colonic LP of sirC versus nsirC within each treatment group. Highlighted genes denote the most significantly upregulated and downregulated transcripts in each comparison. B. Proposed model of FcγR-dependent amplification of checkpoint-specific intestinal inflammation. Anti-PD-1 and anti-CTLA-4 establish distinct inflammatory programs within the colonic lamina propria. Patient-derived sirC IgG engages FcγR-expressing immune cells and amplifies these pre-existing treatment-associated responses rather than inducing an independent inflammatory program. Myeloid cells, ILC3/LTi-like cells, and CD4+ T-cell subsets represent major cytokine-producing compartments, with treatment-specific differences in the relative contribution of these populations. Increased inflammatory signaling may promote leukocyte recruitment and formation of organized inflammatory aggregates/TLS-like structures. The contribution of luminal or microbiota-derived antigenic stimulation remains to be determined.

Together, these data support a model in which patient-derived sirC IgG does not independently initiate irC but instead lowers the threshold for checkpoint-driven colonic inflammation. In both ICI settings, Th1/Th17 and cytotoxic T-cell activation was accompanied by macrophage production of IL-1β and IL-23, IL23R-restricted

## DISCUSSION

Our study identifies a baseline humoral state associated with irC severity and shows that patient-derived IgG can amplify ICI-associated intestinal inflammation in an FcγR-dependent manner. Baseline serum AAb profiling demonstrated that subsequent sirC was associated with a selective antigen signature rather than nonspecific autoreactivity, whereas transfer of polyclonal IgG from patients with sirC was associated with greater colonic inflammation in hFcγR mice, but only in the context of PD-1 or CTLA-4 blockade. Together, these findings support a model in which humoral immunity does not independently initiate irC but instead conditions the magnitude and tissue architecture of checkpoint-driven inflammation.

A notable feature of the pretreatment AAb landscape was the coexistence of two biologically distinct antigen targets. Patients who later developed sirC retained reactivity against melanoma-associated, cancer-testis, and oncogenic antigens, consistent with prior studies linking humoral anti-tumor immunity to ICI-responsive states. In contrast, patients protected from sirC exhibited relative enrichment of AAb directed against immune- and tissue-regulatory targets. In the discovery cohort, these included chemokine receptors and inflammatory mediators (CXCR2, CXCR4, CXCR6, CCR5, and CCL5) together with regulators of epithelial and immune homeostasis such as SOCS3, SMAD4, and CBFA2T3; the validation cohort provided independent support for a related immune- and mucosal-regulatory axis through BRD2, HRH2, and OVOL1. The enrichment of a chemokine-centered protective axis in nsirC patients is biologically plausible given the central role of chemokine networks in leukocyte trafficking, epithelial injury responses, and intestinal immune homeostasis: disruption of CXCR2 signaling attenuates experimental colitis; SMAD4 constrains intestinal inflammation, whereas CBFA2T3 contributes to epithelial differentiation, regeneration after injury, and protection from colitis.

This interpretation shifts the conceptual framework from one centered solely on enhanced autoreactivity to one that also considers the loss of protective immune-regulatory programs. Notably, the AAb depleted in patients who developed sirC converge on pathways controlling leukocyte trafficking and mucosal homeostasis, raising the possibility that they mark host immune states that limit the colonic inflammatory circuits triggered by checkpoint blockade. However, we identified these associations using defined antigen arrays and should not interpret them as evidence that any individual antibody specificity is functionally responsible for the phenotype observed after polyclonal IgG transfer. The AAb landscape and the functional mouse model are complementary lines of evidence for a humoral-regulatory theme in ICI-mediated GI irAEs, not a validated mechanistic chain. Nevertheless, the findings support the concept that baseline humoral immunity is not merely epiphenomenal but may help shape susceptibility to clinically significant irAEs.

The mouse irC modeling through IgG transfer supported this framework. IgG from patients with sirC potentiated intestinal inflammation in hFcγR mice only with checkpoint blockade, consistent with a requirement for IgG-FcγR engagement within a permissive inflammatory context [22, 23, 50, 51]. Existing murine models of irC largely rely on checkpoint blockade in the context of microbiota perturbation or genetically susceptible hosts and have identified T-cell-dependent inflammatory pathways as central mediators of disease. In contrast, the contribution of patient-derived humoral immunity to irC has remained poorly defined. A humanized FcγR model allows testing whether circulating IgG from patients with irC modifies susceptibility to checkpoint inhibitor-associated intestinal inflammation. Serum volume limited the number of recipient mice per donor, so biological replication in these experiments was assessed primarily at the level of recipient mice within each IgG category (sirC, nsirC, HC), an approach consistent with prior IgG passive-transfer studies in which multiple mice per patient donor were used to establish a phenotype [52]. Because multiple recipient mice received IgG from each donor, the recipient-level analyses provide evidence for a phenotype associated with clinical IgG category but do not establish donor-level independence. Donor-level averaging showed the same direction of effect for every comparison, although statistical significance was not retained after collapsing to the smaller number of independent donors. This limitation was greatest in the anti-CTLA-4 arm, which included only three donors per group; those findings should therefore be considered preliminary.

The resulting irC phenotype was mild and most consistent with early or evolving colon inflammation rather than fulminant disease. We view this as an informative feature of the model, although it limits the model’s ability to capture severe clinical disease, because it argues against circulating IgG as a self-sufficient initiator of tissue injury and instead supports an amplification model in which humoral factors lower the threshold for ICI-induced inflammation. The absence of a significant large-scale compositional shift in longitudinal 16S profiling suggests that the observed phenotype did not require a detectable change in overall bacterial community composition during the 21-day experimental window. This result does not exclude functional, spatial, metabolomic, or later microbiome-dependent effects.

Importantly, findings from the hFcγR model converged with observations in human irC. Analysis of an independent patient biopsy cohort identified a myeloid-rich inflammatory niche characterized by abundant CD68+ cells expressing activating FcγRs, including CD16 and CD64, and an overall activating-to-inhibitory FcγR bias. Because the biopsy cohort included only patients with established irC and did not include normal or ICI-treated non-colitis control mucosa, these studies define the immune composition of irC lesions but do not establish whether the observed FcγR-expressing myeloid infiltrate is specific to irC or represents an extension of baseline colonic myeloid composition. In hFcγR mice, single-cell profiling similarly localized activating FcγR expression primarily to macrophage and monocyte populations, with the strongest activating receptor signature observed in inflammatory (M1-like, TM) macrophage populations. Because scRNA-seq libraries were pooled by condition for technical reasons, we treat this transcriptomic localization as descriptive rather than independently replicated at the single-cell level; however, multiplex imaging in mice injected with the same IgG independently confirmed FcγR protein expression on colonic F4/80+ and CD11b+ myeloid cells, providing an orthogonal, protein-level readout that corroborates the transcriptomic finding and is not subject to the same pooling limitation. Together with the human biopsy data, these findings nominate FcγR-expressing myeloid cells as candidate tissue effectors and provide a cellular framework for the FcγR dependence observed in the model. This interpretation is further supported by Bauché et al [51], who recently demonstrated that disruption of FcγR signaling attenuates ICI-associated intestinal inflammation.

This model also suggests that patient-derived IgG does not impose a uniform inflammatory program across ICIs. Rather, sirC IgG amplified checkpoint-specific immune circuits. Under PD-1 blockade, sirC IgG was associated with inflammatory macrophage expansion, cytokine and stress-response programs, and increased circulating IL-17A and IL-22. In contrast, under CTLA-4 blockade, sirC IgG preferentially promoted remodeling of lymphoid compartments, including T-cell, B-cell, and innate lymphoid populations. This context dependence is biologically plausible given the distinct mechanisms of action of CTLA-4 and PD-1 blockade: CTLA-4 acts predominantly during T-cell priming [53], whereas PD-1 regulates effector responses in previously activated peripheral T cells [9]. The checkpoint antibodies themselves also differ in species of origin and isotype - anti-CTLA-4 (Syrian hamster IgG2b, clone 9H10) versus anti-PD-1 (rat IgG2a, clone RMP1-14) - and so may engage the human FcγRs expressed in this model unequally: systematic profiling of cross-species IgG-FcγR interactions indicates that rat IgG2a binds human FcγRs poorly, whereas hamster IgG shows detectable engagement of the activating receptor hFcγRIIA [54]. We cannot exclude that this asymmetry contributes some FcγR-mediated signal specific to the anti-CTLA-4 arm, independent of CTLA-4 blockade itself; Fc-silenced checkpoint antibody variants would be needed to formally dissociate target-driven from Fc-driven contributions to the treatment-specific immune architecture we describe. Within this framework, pre-existing IgG AAb is best viewed not as a dominant upstream driver, but as an amplifier superimposed upon checkpoint-defined inflammatory states.

Notably, the immune programs observed in hFcγR mice recapitulated several key features reported in human irC. Recent single-cell studies have identified expansion of tissue-resident and tissue-adapted lymphocyte populations, Th1/Th17-associated inflammatory programs, IFNγ-producing effector cells, activated mononuclear phagocytes, and dysfunctional regulatory networks within inflamed colonic mucosa [55].

Consistent with these observations, sirC IgG-exposed mice demonstrated Th1/Th17 polarization, emergence of IFNγ-associated Vδ4+ and NKT populations, and expansion of innate-like (ILT) CD8^low^ Ikzf2^high^, Itgae^high^, Itga1^high^ populations. Elevated IL-17A and IL-22 levels were accompanied by expansion of ILC3 populations, supporting activation of mucosal inflammatory pathways implicated in both human irC and inflammatory bowel disease [47, 56]. Collectively, these findings suggest that FcγR-dependent humoral mechanisms converge upon inflammatory circuits already known to drive checkpoint-associated intestinal pathology

Recent studies suggest that chronic stimulation and checkpoint blockade can induce substantial transcriptional and functional plasticity in Tregs, including acquisition of tissue-remodeling or inflammatory features [57]. In our model, we observed an expansion of activated, tissue-adapted Tregs in the inflamed colon, particularly in anti-CTLA-4-treated mice receiving sirC IgG. These Tregs retained classical suppressive markers (*Il10* and *Ctla4)* but also expressed genes associated with activation and tissue adaptation (*Il1rl1* (*ST2*), *Tnfrsf4* (*OX40*), *Tnfrsf9* (*4-1BB*), *Areg*, *Itgav*, *Cpm*, *Zeb2*, and *Dusp4)*. This phenotype is consistent with Treg remodeling in an inflammatory, microbially influenced, cytokine-rich environment. Although our data do not establish loss of suppressive function, they support convergence of antibody-amplified and T cell-driven inflammatory circuits in irC.

A notable intersection between the two checkpoint settings was the emergence of an IL-22-associated mucosal remodeling program. Single-cell analysis localized *Il22* predominantly to ILC compartments, whereas *Il1b* mapped to monocyte/macrophage populations and *Il17a* to CD4 T-cell subsets, revealing a multicompartment inflammatory network spanning innate, adaptive, and tissue-organizing responses. Although our data do not establish a causal role for IL-22, the recurrent appearance of this pathway across both ICI contexts suggests that it may represent a common downstream feature of IgG-amplified intestinal inflammation despite upstream differences between PD-1 and CTLA-4 blockade. Likewise, several populations enriched in our model, including ILC1/NK-like cells, innate-like T cells, and Th1/Th17-like lymphocytes, expressed Ifng and cytotoxic effector genes, closely paralleling the IFNγ-driven programs described in both murine and human irC [8, 10, 58].

In summary, our findings support a model in which pretreatment humoral immunity influences susceptibility to intestinal irAEs by lowering the threshold for checkpoint-driven tissue inflammation rather than independently initiating disease. Patient-derived IgG was associated with FcγR-dependent reshaping of checkpoint-specific inflammatory responses involving myeloid, lymphoid, and mucosal-remodeling programs. The AAb profiling, IgG-transfer experiments, and single-cell and imaging analyses provide complementary evidence for this model, but do not establish a linear antigen-specific mechanism. These findings position the IgG-FcγR axis as a candidate biomarker and mechanistic pathway. FcRn antagonism provides a clinically relevant strategy [59] [60] for testing whether transient reduction of circulating IgG can mitigate IgG-amplified irC.

## Author Contributions

**Project concept and design:** I. Voloshyna, S. Sandigursky, I. Osman, Y. Patskovsky, and M. Krogsgaard

**Clinical evaluation, pathology evaluation, and selection of melanoma patients for the experiments:** F. Fa’ak, E. C. Bayraktar, and I. Osman

**Administrative or material support (specimens’ inventory and retrieval from storage, reporting or organizing data, constructing databases):** I. Voloshyna, S. Idga, C. Ng, and A. V. Lopez.

**Experiments execution and acquisition of data:** I. Voloshyna, S. Sandigursky, M. Ibrahim, A.V. Lopez, S. Idga, C. Ng, A. Zhurova, J. Mastroianni, E. Tardio, and R. Freih

**Data analysis and interpretation, figure preparation:** C. Sreenivasaiah, Y. Hao, I. Voloshyna, Y. Patskovsky, S. Sandigursky, P. Mishra, Alireza Khodadadi-Jamayran, M. Ibrahim, S. Idga, A. Zhurova, R. Freih, G. Silverman, and M. Krogsgaard

**Writing, review, and/or revision of the manuscript:** I. Voloshyna, Y. Patskovsky, S. Sandigursky, M. Ibrahim, J. Mehnert, G. Silverman, I. Osman, and M. Krogsgaard.

All authors have reviewed the manuscript and approved the final manuscript for publication.

## Acknowledgments

High-throughput quantification of autoantibodies and related data analysis were performed with technical support from Sengenics Corporation LLC. Single-cell RNA sequencing and 16S rRNA library preparation and sequencing were performed at NYU Grossman School of Medicine Genome Technology Center. OPAL multiplex fluorescent staining, imaging, and data analysis were performed in the NYU Grossman School of Medicine Experimental Pathology Research Laboratory. We thank Valeria Mezzano (Experimental Pathology Research Laboratory) for her help with OPAL multiplex imaging analysis. We thank Duane Moogk (McMaster University) and Jean M. Etersque (Grants Writer at the Laura and Isaac Perlmutter Cancer Center) for critically reviewing the manuscript and providing valuable suggestions that improved its clarity and presentation.

## Data Availability

All data generated in this study are available from the Gene Expression Omnibus (GEO) under accession number TBD. Single-cell RNA sequencing and multiplex imaging datasets are also stored in the NYU Grossman School of Medicine Core Facility Database. The corresponding author (M.K.) will evaluate requests for raw data and, where appropriate, fulfill them under a material transfer agreement.

## Ethics Statement

This study complied with all relevant ethical regulations. All patient samples used in this study were collected under protocols approved by the respective institutional regulatory committees (see the Methods section) and were performed as part of the standard of care. We obtained written informed consent from all patients. We performed *in vivo* experiments in compliance with the reference protocol (IA16-00618) approved by the NYU Institutional Animal Care and Use Committee (IACUC).

## METHODS

### Patient Enrollment, Serum, and Biopsy Collection

We included patients with melanoma treated at NYU Langone Health between 2012 and 2020 under the Interdisciplinary Melanoma Cooperative Group (IMCG) biospecimen protocol approved by the Institutional Review Board (IRB 10362). Under this protocol, we prospectively banked biospecimens for research after written informed consent for collection of clinical data and specimens at enrollment. The discovery cohort comprised 149 patients treated with anti–PD-1 therapy in the CheckMate 915 adjuvant clinical trial, including 31 patients who subsequently developed severe irC (sirC; **Table S1**) and 118 patients with non-severe irC (nsirC; **Table S1**). We used pretreatment serum samples from this cohort for baseline AAb profiling. An independent experimental cohort for IgG purification and mechanistic studies included 18 patients with stage III/IV melanoma treated with anti-PD-1 (n = 12) or anti-CTLA-4 (n = 6). We classified patients by peak colitis severity according to CTCAE criteria as non-severe immune-related colitis (nsirC; grades 1–2) or severe immune-related colitis (sirC; grades 3–4; **Table S2**). Healthy volunteers (n = 9) served as an additional comparator group. To minimize preanalytical variability, we collected, processed, and stored blood samples according to standardized NYU IMCG protocols. We collected whole blood in Becton Dickinson Vacutainer SST tubes and processed it within 90 minutes of collection by centrifuging at 2,500 rpm for 10 minutes at room temperature. We aliquoted serum into 1.8-mL cryovials (1 mL), stored it at −80°C, and thawed it once for proteomic array analysis.

To define the histologic and immune context of human irC for comparison with hFcγR mouse studies, formalin-fixed, paraffin-embedded (FFPE) colon biopsy specimens were obtained from patients who developed irC through the NYU Center for Biospecimen Research and Development. This cohort (n=10) included biopsies from 2 patients with enterocolitis following combined ipilimumab plus nivolumab, and 8 patients with colitis following anti-PD-1 monotherapy (nivolumab or pembrolizumab). We had no access to non-inflamed normal colon biopsies and ICI-treated biopsies from patients without irC, therefore, they were not included in this analysis. The enterocolitis specimens were from patients whose sera were also used for IgG isolation and mechanistic modeling in hFcγR mice, collected before treatment. Colon biopsies were obtained from melanoma patients with clinically diagnosed irC spanning mild, chronic, and enterocolitis phenotypes. After centralized pathology review, we used all biopsies for histopathologic and multiplex imaging analyses (**Fig. 1A**). We classified histology based on established features, including crypt architectural distortion, lamina propria inflammation, intraepithelial neutrophils, crypt abscesses, epithelial apoptosis, and ulceration (**Fig. S3**). The median interval between ICI initiation and biopsy was 24 weeks.

### Isolation and Purification of Human Serum IgG

Polyclonal IgG was isolated from patient serum using protein G agarose (Thermo Scientific; Cat# 22852)[61]. Serum was diluted in sterile cold binding buffer to a final composition of 0.1 M sodium phosphate and 0.15 M NaCl (pH 7.4). Particulate material was removed by centrifugation or filtration through a 0.8 -µm filter. Protein G agarose was equilibrated with binding buffer and incubated with diluted serum at a 1:1 resin-to-serum ratio for 1 h at room temperature with gentle mixing. Following binding, the resin was packed into a gravity-flow column and washed with binding buffer. Bound IgG was eluted with an acidic buffer, and individual fractions were collected into tubes preloaded with concentrated neutralization buffer to achieve near-neutral pH immediately upon collection. Eluted IgG was quantified and transferred into Slide-A-Lyzer dialysis cassettes (10,000 MWCO; Thermo Scientific, Cat# 66456) for overnight dialysis against 1 L of 1× PBS at 4°C. We concentrated IgG using Amicon Ultra-15 centrifugal filters with a 30-kDa molecular-weight cutoff (Millipore, Cat. No. UFC903024), passed it through a 0.22-µm sterile filter, and adjusted it to 25 mg/mL in sterile PBS. We evaluated IgG purity by reducing and nonreducing SDS-PAGE with Coomassie staining. For *in vivo* experiments, mice received 5 mg human IgG in 200 µL sterile PBS by intraperitoneal injection.

### AAb Profiling

Baseline serum samples from the discovery and experimental cohorts were profiled using KREX™ protein microarrays (former Sengenics Co., LLC), which quantify IgG AAb reactivity against >1,800 correctly folded human proteins. We included five pooled normal-serum internal controls across runs as technical quality-control replicates. To assess normalization performance, we evaluated concordance among internal control samples before downstream analysis. We normalized raw signal intensities to reduce systematic technical variation using LOESS and ComBat-based batch correction [62]. We analyzed differential AAb titers across array features using the limma package with moderated t-statistics [63]. Prespecified comparisons included sirC versus nsirC, sirC versus healthy controls, and treatment-stratified subgroup analyses, as indicated in the corresponding figure legends. For each antigen, we summarized the effect size as a fold change and assessed significance using model-derived p-values with a Benjamini–Hochberg correction for multiple testing. We used receiver operating characteristic [52] analysis [64] to assess the discriminatory performance of selected antigens, and calculated area under the curve (AUC) values [16]. For visualization and candidate prioritization, antigens were ranked based on adjusted P-value, fold change, and AUC.

Because we used purified donor IgG for passive-transfer and mechanistic studies, we also profiled matched serum and purified IgG samples from the same patients to determine whether antigen-specific reactivities detected in whole serum were retained after IgG isolation. These profiles were highly concordant (Spearman r > 0.9), indicating that the measured AAb titers were predominantly IgG-derived and were preserved in the material used for *in vivo* studies. We then performed differential reactivity analysis to identify antigens associated with sirC, nsirC, or HC groups using a prespecified significance threshold of adjusted P < 0.05.

### Mice Used in the Study

C57BL/6J mice (Strain #000664) were purchased from The Jackson Laboratory. Humanized Fcγ receptor (hFcγR) mice were kindly provided by the laboratory of Dr. Jeffrey Ravetch (Rockefeller University). We generated this strain on a C57BL/6 background as previously described [22]. Briefly, a targeted 95-kb deletion encompassing the murine FcγR α-chain locus on chromosome 1 (Fcgr2b, Fcgr3, and Fcgr4) was combined with deletion of Fcgr1 (FcγRI) on chromosome 4, resulting in complete ablation of endogenous murine FcγR expression. Transgenes encoding human Fcγ receptors (FCGR1A, FCGR2A, FCGR2B, FCGR3A, and FCGR3B) were introduced to reconstitute human FcγR expression *in vivo.* Transnetyx performed genotyping to maintain and validate colonies. We confirmed human FcγR expression by multiplex tissue imaging and scRNA-seq. All animal experiments were performed in accordance with the guidelines of the Institutional Animal Care and Use Committee (IACUC), protocol IA16-00618. Mice were housed in ventilated, autoclaved, pathogen-free cages with unrestricted access to food and water, maintained on a 12-h light/dark cycle at 22 ± 1 °C and 55 ± 10% humidity. All experimental animals were housed within the same facility and room to minimize environmental variability.

### Experimental Design

The current study was designed to evaluate whether patient-derived IgG modulates ICI-associated intestinal inflammation in an FcγR-dependent manner. Purified serum IgG was obtained from melanoma patients in the experimental cohort and grouped according to donor phenotype as sirC (grade 3–4), nsirC (grade 1–2), or healthy control (HC) (**Fig. 2A**). Recipient hFcγR mice received two intraperitoneal injections of purified human IgG (5 mg in 200 μL) on days 0 and 2 (**Fig. 2A**). In selected comparator experiments, WT C57BL/6 mice received the same IgG transfer protocol. Beginning on day 2, mice received anti-PD-1 or anti-CTLA-4 monoclonal antibody, matched to the ICI class received by the corresponding IgG donor, or the appropriate isotype control. We administered checkpoint antibodies intraperitoneally on days 2, 6, and 8. Additional control groups included untreated hFcγR mice and hFcγR mice receiving IgG without checkpoint blockade. We monitored mice from day 2 until study completion for weight change, fur changes, skin lesions, postural abnormalities, and rectal bleeding. We collected fecal pellets on days 0, 6, 13, and 20 for lipocalin-2 measurement and microbiome analysis. Mice were euthanized on day 21, and blood was collected by cardiac puncture for serum isolation.

We used 2–6 recipient mice per donor IgG preparation, an approach consistent with prior studies using individual patient-derived IgG in an experimental setting [52]. For single-cell RNA-seq and OPAL multiplex imaging, we randomized donor-matched recipient pairs so one mouse per pair was allocated to each assay, ensuring these two readouts came from independent mice within each donor. For histologic scoring, multiple recipient mice per donor were evaluated; to account for non-independence among mice receiving IgG from the same donor, histology scores were additionally averaged across all recipient mice for each IgG donor to generate a donor-level score, and comparisons were repeated using the IgG donor as the unit of biological replication (**Supplementary Fig. S5A**; **Supplementary Table S1**).

All mouse experiments were performed with at least 3 biological replicates unless otherwise indicated. Data were analyzed and plotted using GraphPad Prism v10.4.0. For comparisons between two groups, a two-tailed Mann-Whitney test was used, except where otherwise specified. For repeated-measures comparisons over time (e.g., fecal lipocalin-2), we used two-way ANOVA with Tukey’s post-hoc test for pairwise comparisons. For comparisons across a multi-analyte panel (e.g., serum cytokine profiling), we used the Benjamini-Hochberg procedure to correct for multiple comparisons. P < 0.05 (or q < 0.05 for Benjamini-Hochberg–corrected comparisons) was considered statistically significant unless otherwise specified.

### Treatment with Immune Checkpoint Inhibitors

Mice received intraperitoneal injections of anti-mouse CTLA-4 antibody (clone 9H10; 200 µg per dose; BioXCell, Cat. No. BP0131), anti-mouse PD-1 antibody (clone RMP1-14; 200 µg per dose; BioXCell, Cat. No. BP0146), or the corresponding species- and isotype-matched control antibodies. Because clone 9H10 is a Syrian hamster-derived antibody, we used polyclonal Syrian hamster IgG as its control (BioXCell, Cat. No. BP0087). Because clone RMP1-14 is a rat IgG2a antibody, rat IgG2a was used as its matched control (BioXCell, Cat. No. BP0089).

### Histology and Histopathological Evaluation

Whole colons were collected at day 21 and processed using the Swiss roll technique [65], with the distal end rolled inward and the luminal surface facing outward. Tissues were paraffin-embedded, sectioned at 5 µm, and processed by the Experimental Pathology Research Laboratory at NYU Langone. Sections were mounted on glass microscope slides and stained with hematoxylin and eosin (H&E) using standard procedures. Two pathologists independently evaluated colonic histopathology, blinded to treatment group. The following features were scored: goblet-cell loss (0, absent; 1, present), submucosal inflammatory-cell infiltration (0, absent; 1, present), mucosal leukocyte infiltration (0, absent; 1, mild; 2, severe), and lymphocyte aggregates (0, absent; 1, 1–3 foci; 2, >3 foci). We summed individual scores to generate a cumulative histopathology score using a grading system modified from a previously published method [38] **(Supplementary Table S3**).

### Serum Cytokine and ALT Analysis

We retrospectively analyzed serum samples from melanoma patients and mice to quantify cytokines and assess liver function via serum alanine aminotransferase (ALT) levels [41]. Cytokine profiling included IL-17A, IL- 12(p70), IL-6, IL-1β, IL-4, TNFα, IL-22, IL-10, and IFNγ, measured using the MILLIPLEX Mouse TH17 Magnetic Bead customized testing panel (Millipore; Cat# MTH17MAG-47K-09) for mice, the MILLIPLEX Human TH17 Magnetic Bead Panel (Millipore; Cat# HTH17MAG-14K) for human samples, and the Luminex 200 system (Luminex), according to the manufacturer’s instructions. We calculated cytokine concentrations (pg/mL) from standard curves using mean fluorescence intensity, normalized for batch and plate effects; values below the detection threshold were excluded. To assess liver function in mice, we measured serum ALT levels using a Mouse ALT ELISA kit (#M7832). We performed quantitative analysis on 5-fold-diluted serum after sample buffer blocking, incubation with biotinylated ALT antibody at 37°C for 20 minutes, and incubation with HRP-streptavidin conjugate at 37°C for 10 minutes. After washing, plates were developed with TMB substrate and read at 450 nm. ALT concentrations were determined using standard curves.

### Mouse Fecal Lipocalin 2 (LCN2)

We measured fecal lipocalin 2 (LCN2), a sensitive and non-invasive biomarker of intestinal inflammation, on days 0, 6, 13, and 20 per the study design. We measured LCN2 levels using an ELISA kit according to the manufacturer’s protocol (R&D Systems, Cat# DY1857). Briefly, we diluted supernatants from fecal samples 50- fold using a kit-recommended reagent diluent (1% BSA in PBS). We read the plates at 450 nm with correction at 570 nm. We measured fecal sample weights before the assay. Fecal LCN2 levels were calculated as the concentration per 1 mg of solids.

### Isolation of Lamina Propria Mononuclear Cells (LPMCs) and 3′ scRNA-seq Library Preparation

We isolated LPMCs from mouse colonic tissue using previously established protocols [65, 66]. Briefly, visible fat and Peyer’s patches were removed, and intestines were flushed with phosphate-buffered saline (PBS; without calcium or magnesium) at room temperature to remove luminal contents. Tissues were opened longitudinally, washed thoroughly to remove mucus, cut into 1-cm pieces, and incubated in HBSS (Gibco, Cat# 14170088; without calcium or magnesium) supplemented with 10% FBS and 2 mmol/L EDTA to remove epithelial cells. We then digested the tissue using the Miltenyi Biotec Lamina Propria Dissociation Kit, following the manufacturer’s instructions. We pelleted the released cells by centrifugation and washed them in RPMI 1640 supplemented with 10% FBS, 100 U/mL penicillin, 100 μg/mL streptomycin, and 2 mmol/L L-glutamine. Recovered cells were stained with TruStain FcX (anti-mouse CD16/32, BioLegend, clone 93), DAPI viability dye (BioLegend, Cat# 423106), and CD45-PerCP/Cy5.5 (BioLegend, clone 30-F11). Viable CD45+ cells were sorted on a FACSAria IIu SORP cell sorter, washed in PBS containing 0.5% RNase-free BSA, and counted prior to library preparation. For scRNA-seq, we pooled cells within each experimental subgroup before loading onto the 10x Genomics Chromium platform, so each library represented one subgroup sample from 4 mice. Subgroups included sirC, nsirC, HC, and sirC + isotype ICI within each checkpoint-treatment experiment. We attempted cell hashing to enable sample demultiplexing, but its technical performance was inadequate to reliably assign individual mouse origins; accordingly, we conducted downstream analyses at the pooled subgroup level. Equal numbers of viable CD45+ cells from each pooled subgroup sample (up to 100,000 cells) were loaded into a single-cell chip together with reverse transcription reagents and gel beads according to the manufacturer’s instructions. Single-cell RNA-seq libraries were prepared using the 10x Genomics Chromium Single Cell 3′ Reagent Kits v3.1, including the GEM, Library & Gel Bead Kit v3.1 (PN-1000121), Chip G Single Cell Kit (PN-1000120), and Single Index Kit T Set A (PN-1000213). Libraries were sequenced on an Illumina NovaSeq 6000 platform using paired-end reads (28 cycles for Read 1, 8 cycles for the i7 index, and 91 cycles for Read 2), with one lane per sample.

### Multiplex Immunofluorescence of FFPE colon

Five-micron paraffin-embedded sections were stained either with hematoxylin and eosin or with Akoya Biosciences® Opal™ multiplex automation kit reagents (Leica Cat #ARD1001EA) on a Leica BondRX® autostainer, according to the manufacturer’s instructions. Briefly, slides were subjected to heat-induced epitope retrieval (Leica Biosystems epitope retrieval 2 solution ER2, EDTA-based, pH9, Cat. AR9640), followed by a protein block (ARD1001EA, Akoya Biosciences), sequential primary and secondary antibody incubations, and tyramide signal amplification with covalent linkage of opal fluorophores (Op480, 520, 570, 620, 690, 780; Akoya Cat #FP1488001KT or Op690 Akoya Cat #FP1497001KT) to the underlying antigen. We optimized IHC labeling as previously described [67, 68]. We used two panels to characterize human colon immune cells (**Supplementary Table S4**) and mouse colon immune cells (**Supplementary Table S5**). Panel 1 includes F4/80- b (CD68a in humans), Ly6g (Arg1a in humans), CD11-b, CD4, CD8, and FOXP3. Panel 2 shows the staining for CD19 (CD11b), CD3, CD64, CD32b, and CD16. After each staining round, we removed primary and secondary antibodies via epitope retrieval. Slides were counterstained with spectral DAPI (Akoya Biosciences, FP1490) and mounted with ProLong Gold Antifade (ThermoFisher Scientific, P36935). Semi-automated image acquisition was performed on a Vectra® Polaris multispectral imaging system (Akoya PhenoImages HT) in Motif mode. After whole slide scanning at 20×, the tissue was manually outlined to select fields for spectral unmixing using InForm® version 2.4.11 software from Akoya Biosciences.

### Gut Microbiota Analysis with 16S rRNA Gene Sequencing

We freshly harvested feces at critical time points (D0, D13, and D21) and froze them at -80 °C until DNA isolation. We extracted microbial genomic DNA from 20 mg (on average) of feces using a DNeasy PowerSoil® Pro Kit (Qiagen) according to the manufacturer’s instructions. After elution, we determined sample yield and purity using a NanoDrop spectrophotometer (Thermo Scientific NanoDrop 2000). We stored the samples at 20 °C until gene sequencing. We assessed the fecal microbiota using 16S rRNA sequencing. We prepared the amplicon library on an automated platform (Biomek 4000) using a custom liquid-handling method. We amplified the V4 region of the 16S rRNA gene using barcoded primers (Parada et al., 2016). Each unique barcoded amplicon was generated in pairs of 25 µL reactions under the following reaction conditions: 11 µL PCR-grade H2O, 10 µL HotMasterMix (QuantaBio), 2 µL of barcoded forward and reverse primers (5uM), and 2 µL of 100× diluted template DNA. reactions were run on a C1000 Touch Thermal Cycler (Bio-Rad) under the following cycling conditions: initial denaturation at 94°C for 3 min, followed by 35 cycles of denaturation at 94°C for 45 s, annealing at 50°C for 1 min, and extension at 72 °C for 90 s, with a final extension of 10 min at 72°C. We quantified amplicons using an Agilent 2200 TapeStation system and pooled them. Purification was performed using Ampure XT (Beckman Coulter), according to the manufacturer’s instructions, and the final concentration was determined using Qubit (Invitrogen). We performed sequencing as paired-end, 250 cycles on a MiSeq (Illumina).

### Processing Data of 16S rRNA Sequences

We demultiplexed and analyzed the paired-end FASTQ sequences using QIIME 2 version 2–2020.2 (https://qiime2.org). We used the DADA2 algorithm [69] in QIIME2 for error modeling and FASTQ filtering, with parameters p-trunc-len-f 150 --p-trunc-len-r 150, for both the anti-CTLA-4 and anti-PD-1 experimental groups. These parameters specified the overlap between forward and reverse reads in DADA2. We performed taxonomic classification using the QIIME2 feature classifier classification consensus search plugin trained on the Silva 132 database [70]. We calculated alpha and beta diversities using a rarefied sampling depth of 1000. OTU tables were exported as matrices for CTLA4 and PD1 using qiime2R (https://github.com/jbisanz/qiime2R). We converted these absolute counts to relative abundances by dividing each count by the total number of counts across all phylogenetic species in its treatment group. We generated a relative abundance plot using ggplot2 (v3.3.5). We evaluated beta diversity using principal coordinates analysis (PCoA) of unweighted UniFrac distances. Shannon’s index was calculated using “alpha group significance in QIIME2 to extract information on the distribution of microbiota within samples. We used t-tests to compare means between treatment groups, with alpha set at 0.05 and the Bonferroni test to correct for multiple comparisons.

### Processing of Single-Cell Transcriptome Data and Differential Expression Analysis

For each experimental subgroup, we pooled cells from four mice for single-cell RNA-sequencing (scRNA-seq) analysis. After confirming the integrity of the cDNA, quality of the libraries, number of cells sequenced and mean number of reads per cell as quality control, we used the Cellranger package to map the reads and generate gene-cell matrices [58]. We then performed quality control to calculate the number of genes, UMIs, and the proportion of mitochondrial genes for each cell, and filtered out cells with a low number of covered genes (gene-count < 500) and high mitochondrial counts (mt-genes > 0.2). We then normalized the matrix by library size. A general statistical test was then performed to calculate gene dispersion, base mean and cell coverage to use to build a gene model for performing Principal Component Analysis (PCA). Genes with high coverage and high dispersion (dispersion > 1.5) were chosen (2000 genes) to perform PCA and Mutual Nearest Neighbors (MNN) batch alignment using the iCellR package (v1.5.5) (https://CRAN.R-project.org/package=iCellR). T-distributed Stochastic Neighbor Embedding (t-SNE), Uniform Manifold Approximation and Projection (UMAP).

We performed initial cell-type annotation using SingleR (v1.0.6), with the Human Primary Cell Atlas and the Immunological Genome Project (ImmGen) reference datasets [37]. We further refined annotations manually based on canonical marker genes and differential expression patterns. We aligned single-cell RNA sequencing to a combined mouse (mm10) and human (hg38) reference genome. SingleR calculated cluster-level expression profiles using a correlation-based iterative algorithm. We further refined annotations manually based on canonical marker genes and differential expression patterns. Mixed-cell clusters (e.g., CD4/CD8 double-positive T cells or clusters co-expressing B and T cell markers) were split into new clusters based on their transcriptional profiles.

### Spatial Analysis

We exported data from InForm and analyzed them in R to assess cell-subset density and spatial relationships between distinct cell subtypes. To map each cell’s location, we performed a proximity (Nearest-Neighbor) analysis of colonic cells with CD4+ and CD8+ phenotypes and assessed differences in spatial distribution between tox and no-tox mice. We merged and consolidated the cell segmentation files exported by InForm for each phenotype and mouse into a single table in R using the phenoptr package (v0.2.10). The per-cell nearest-neighbor distances for each phenotype were calculated using the find_nearest_distance function. We examined and visualized distances using customized R scripts.

---

**Table S1.**
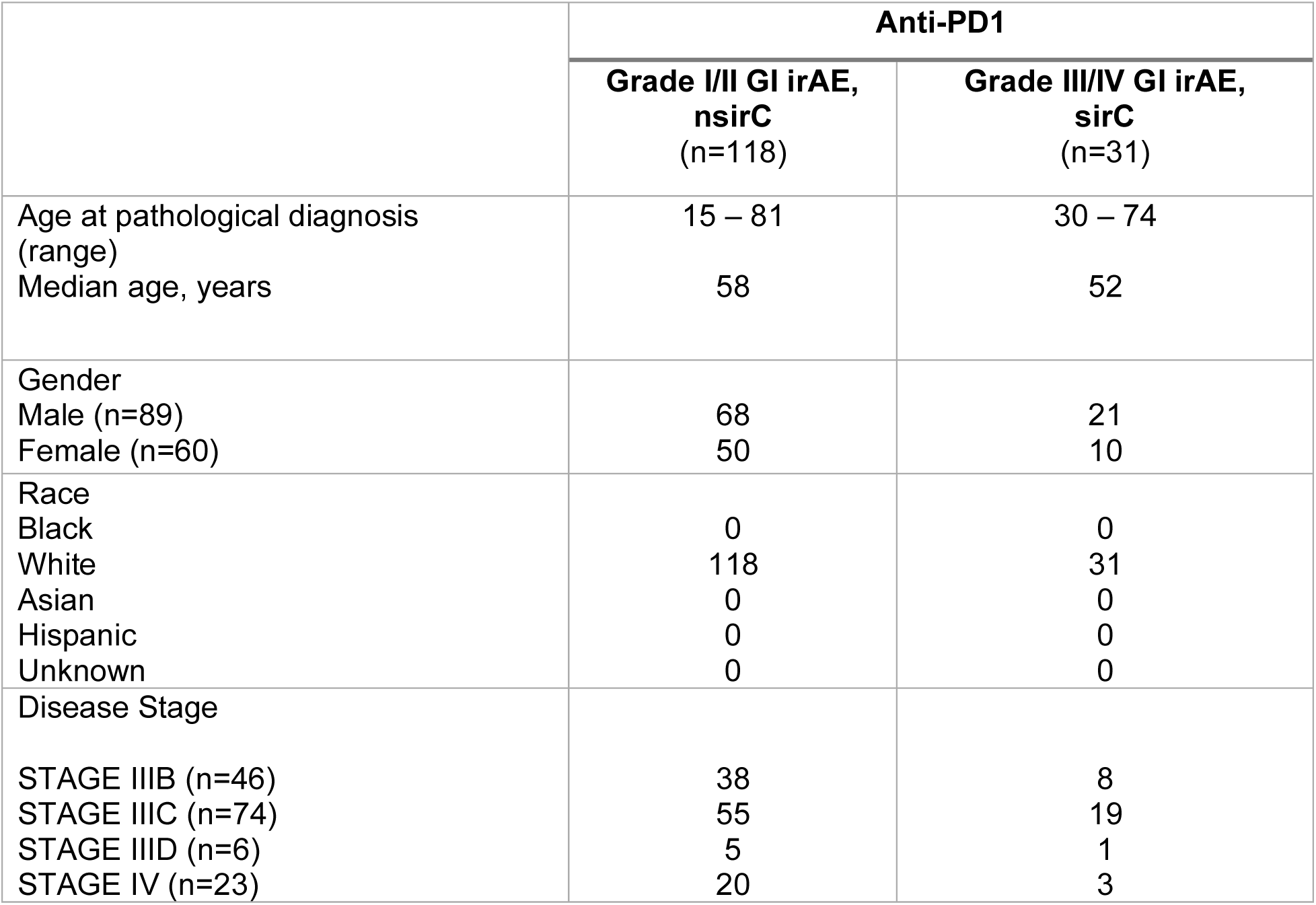
Demographic and clinical characteristics of melanoma patients from the Checkmate 915 trial served as the discovery cohort for AAb screening.

**Table S2.**
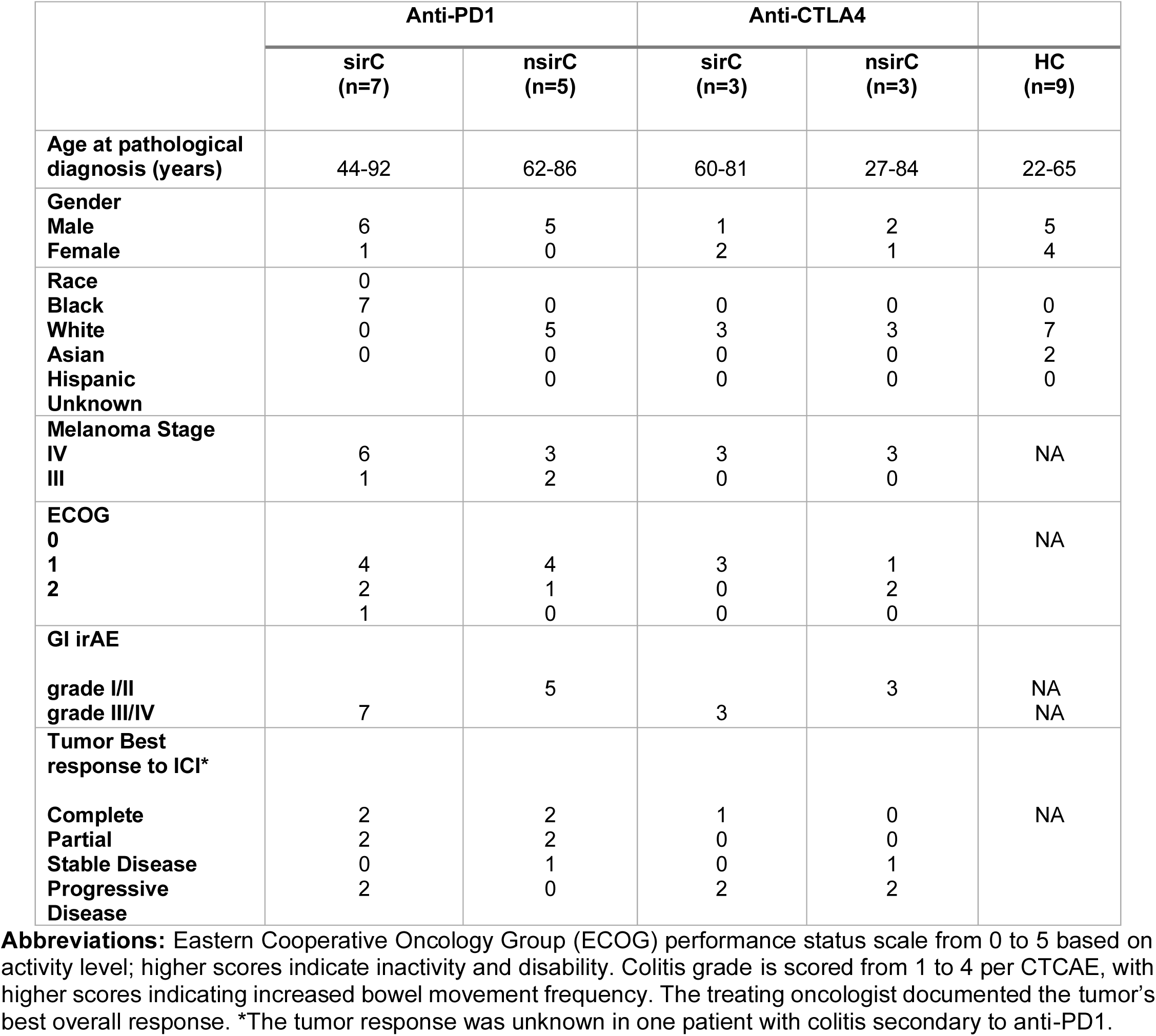
Demographic and clinical characteristics of melanoma patients selected for *in vivo* modeling of irC.

