## Supplementary materials for "Patient-derived IgG Amplifies Fcγ Receptor-Dependent Colonic Inflammation During Immune Checkpoint Blockade"

**Running title: The role of autoantibodies in checkpoint inhibitor-induced colitis**

[1]Laura and Isaac Perlmutter Cancer Center, New York University Grossman School of Medicine, New York, NY 10016, USA.

[2]Department of Pathology, New York University Grossman School of Medicine, New York, NY, 10016, USA

[3]Ronald O. Perelman Department of Dermatology, New York University Grossman School of Medicine, New York, NY, 10016, USA

[4]Applied Bioinformatics Laboratories, New York University Langone Medical Center, New York, NY, 10016, USA

[5]Department of Medicine, New York University Grossman School of Medicine, New York, NY, 10016, USA

**Corresponding author:**
Michelle Krogsgaard, PhD
Associate Professor
Department of Pathology, New York University Grossman School of Medicine, New York, NY, USA.
Laura and Isaac Perlmutter Cancer Center at NYU Langone Health, New York, NY, USA.


**Suuplementary Table S1. Demographic and clinical characteristics of melanoma patients from the Checkmate 915 trial served as the discovery cohort for AAb screening.**

| | **Anti-PD1** | |
|---|---|---|
| | **Grade I/II GI irAE, nsirC (n=118)** | **Grade III/IV GI irAE, sirC (n=31)** |
| Age at pathological diagnosis (range)<br>Median age, years | 15 – 81<br><br>58 | 30 – 74<br><br>52 |
| Gender<br>Male (n=89)<br>Female (n=60) | <br>68<br>50 | <br>21<br>10 |
| Race<br>Black<br>White<br>Asian<br>Hispanic<br>Unknown | <br>0<br>118<br>0<br>0<br>0 | <br>0<br>31<br>0<br>0<br>0 |
| Disease Stage<br><br>STAGE IIIB (n=46)<br>STAGE IIIC (n=74)<br>STAGE IIID (n=6)<br>STAGE IV (n=23) | <br><br>38<br>55<br>5<br>20 | <br><br>8<br>19<br>1<br>3 |

**Table S2. Demographic and clinical characteristics of melanoma patients selected for in vivo modeling of irC (the experimental cohort).**

| | Anti-PD1 | | Anti-CTLA4 | | |
|---|---|---|---|---|---|
| | **sirC (n=7)** | **nsirC (n=5)** | **sirC (n=3)** | **nsirC (n=3)** | **HC (n=9)** |
| Age at pathological diagnosis (years) | 44-92 | 62-86 | 60-81 | 27-84 | 22-65 |
| Gender<br>Male<br>Female | <br>6<br>1 | <br>5<br>0 | <br>1<br>2 | <br>2<br>1 | <br>5<br>4 |
| Race<br>Black<br>White<br>Asian<br>Hispanic<br>Unknown | <br>0<br>7<br>0<br>0 | <br>0<br>5<br>0<br>0 | <br>0<br>3<br>0<br>0 | <br>0<br>3<br>0<br>0 | <br>0<br>7<br>2<br>0 |
| Melanoma Stage<br>IV<br>III | <br>6<br>1 | <br>3<br>2 | <br>3<br>0 | <br>3<br>0 | <br>NA |
| ECOG<br>0<br>1<br>2 | <br><br>4<br>2<br>1 | <br><br>4<br>1<br>0 | <br><br>3<br>0<br>0 | <br><br>1<br>2<br>0 | <br>NA |
| GI irAE<br><br>grade I/II<br>grade III/IV | <br><br><br>7 | <br><br>5 | <br><br><br>3 | <br><br>3 | <br><br>NA<br>NA |
| Tumor Best response to ICI*<br><br>Complete<br>Partial<br>Stable Disease<br>Progressive Disease | <br><br>2<br>2<br>0<br>2 | <br><br>2<br>2<br>1<br>0 | <br><br>1<br>0<br>0<br>2 | <br><br>0<br>0<br>1<br>2 | <br><br>NA |

**Abbreviations:** Eastern Cooperative Oncology Group (ECOG) performance status scale from 0 to 5 based on activity level; higher scores indicate inactivity and disability. Colitis grade is scored from 1 to 4 per CTCAE, with higher scores indicating increased bowel movement frequency. The treating oncologist documented the tumor's best overall response. *The tumor response was unknown in one patient with colitis secondary to anti-PD1.

**Supplementary Table S3. The scoring criteria for the assessment of leukocyte infiltration in mouse colon.**

| Score | Goblet cell loss | Sub-mucosal leukocyte infiltration | Degree of mucosal leukocyte infiltration | Number of lymphocyte aggregates |
|---|---|---|---|---|
| **0** | Absent | Absent | Absent | 0 |
| **1** | Present | Present | Mild | 1-3 |
| **2** | Present | Present | Severe | >3 |

**Supplementary Table S4. Antibodies used for multiplex immunofluorescence on FFPE human colon biopsies.**

**Panel 1:**

| | | | | | | | |
|---|---|---|---|---|---|---|---|
| **CD68** | Dako | Mouse | KP1 | M081401-2 | 1:200 | 570, Akoya, FP1488001KT | No retrieval |
| **CD11b** | Novus | Rabbit | Polyclonal | NB110-89747 | 1:800 | 620, Akoya, FP1495001KT | ER2-20 min |
| **CD4** | Abcam | Rabbit | EPR6855 | ab133616 | 1:200 | 520, Akoya, FP1487001KT | ER2-20 min |
| **CD8** | Dako | Mouse | C8/144B | M710301-2 | 1:200 | 690, Akoya, FP1497001KT | ER2-20 min |
| **Foxp3** | eBioscience | Mouse | D6O8R | 14-4777-82 | 1:75 | 780, Akoya, FP1501001KT | ER2-20 min |
| **Arg1b** | GeneTex | Rabbit | Polyclonal | GTX109242 | 1:1000 | 480, Akoya FP1500001KT | ER1-20 min |

**Panel 2:**

| | | | | | | | |
|---|---|---|---|---|---|---|---|
| **CD16** | Invitrogen | Rabbit | Polyclonal | PA5-80622 | 1:1000 | 780, Akoya, FP1501001KT | ER2-20 min |
| **CD64** | Invitrogen | Rabbit | Polyclonal | PA5-102382 | 1:3000 | 570, Akoya, FP1488001KT | ER1-20 min |
| **CD32b** | Invitrogen | Rabbit | XF3604156 | MA5-35980 | 1:3000 | 690, Akoya, FP1497001KT | ER1-20 min |
| **CD68** | Dako | Mouse | KP1 | M081401-2 | 1:100 | 480, Akoya FP1500001KT | ER2-20 min |
| **CD11b** | Novus | Rabbit | Polyclonal | NB110-89747 | 1:800 | 620, Akoya, FP1495001KT | ER2-20 min |

**Supplementary Table S5. Antibodies used for multiplex immunofluorescence analysis of FFPE mouse colon sections.**

**Panel 1:**

| | | | | | | | |
|---|---|---|---|---|---|---|---|
| **F4/80** | Thermo Fisher | A3-1 | MA1-91124 | 1:100 | Rat HRP - Polymer 1-step (Mouse adsorbed) Biocare, BRR4016 | 480, Akoya FP1500001KT | No retrieval |
| **CD11b** | Novus | polyclonal | NB110-89474 | 1:2000 | Rabbit-on-Rodent HRP-Polymer, Biocare RMR622 | 620, Akoya, FP1495001KT | ER2-20 min |
| **CD4** | CST | D7D2Z | 25229S | 1:500 | Rabbit-on-Rodent HRP-Polymer, Biocare RMR622 | 690, Akoya, FP1497001KT | ER2-20 min |
| **CD8** | CST | D4W2Z | 98941S | 1:300 | Rabbit-on-Rodent HRP-Polymer, Biocare RMR622 | 570, Akoya, FP1488001KT | ER2-20 min |
| **Foxp3** | CST | D6O8R | 12653S | 1:3000 | Rabbit-on-Rodent HRP-Polymer, Biocare RMR622 | 780, Akoya, FP1501001KT | ER2-20 min |
| **Ly6g** | BD | 1A8(RUO) | 551459 | 1:400 | Rabbit-on-Rodent HRP - Polymer 1-step (Mouse adsorbed) Biocare, BRR4016 | 520, Akoya, FP1487001KT | ER1-60 min |

**Panel 2:**

| | | | | | | | |
|---|---|---|---|---|---|---|---|
| **CD3** | CST | E4T1B | 78588S | 1:600 | Rabbit-on-Rodent HRP-Polymer, Biocare RMR622 | 690, Akoya, FP1497001KT | ER2-20 min |
| **CD64** | Invitrogen | Polyclonal | PA5-102382 | 1:3000 | Rabbit-on-Rodent HRP-Polymer, Biocare RMR622 | 480, Akoya FP1500001KT | ER1-20 min |
| **CD32b** | Invitrogen | XF3604156 | MA5-35980 | 1:2000 | Rabbit-on-Rodent HRP-Polymer, Biocare RMR622 | 570, Akoya, FP1488001KT | ER1-20 min |
| **CD19** | CST | D4V4B | 90176S | 1:400 | Rabbit-on-Rodent HRP-Polymer, Biocare RMR622 | 780, Akoya, FP1501001KT | ER2-20 min |
| **CD11b** | Novus | Polyclonal | NB110-89747 | 1:2000 | Rabbit-on-Rodent HRP-Polymer, Biocare RMR622 | 620, Akoya, FP1495001KT | ER2-20 min |

**Figure legends.**

**Supplementary Figure S1. Baseline serum AAb profiling in the CheckMate 915 discovery cohort identifies candidate antigens associated with irC severity. A.** Heatmap showing 40 most differentially enriched AAb targets identified in melanoma patients with sirC compared with nsirC patients. Rows represent autoantigens and columns represent individual patient serum samples. Color intensity reflects normalized AAb titers, with red indicating higher and blue indicating lower relative abundance. **B.** Distribution of serum AAb titers to selected candidate autoantigens at the individual patient level. Each data point represents a single melanoma patient and corresponding normalized AAb titer. Multiple unpaired two-tailed *t*-tests were used to analyze the data.

**Supplementary Figure S2. Serum AAb profiling in experimental melanoma cohort used for IgG isolation and *in vivo* modeling of irC. A.** Heatmaps of serum log2-normalized AAb titers of melanoma-associated antigens in the independent donor cohort (n = 18), selected for IgG purification and mechanistic studies. Rows represent antigens and columns represent individual patient serum samples. **B.** Top-ranked AAb targets associated with sirC in anti-PD-1 (aPD-1) and anti-CTLA-4 (aCTLA-4) treated patients from the *in vivo* modeling cohort, ranked by area under the receiver operating characteristic curve (AUC). Data are plotted as log2-normalized signal intensity across sirC, nsirC, and HC groups. Corresponding fold-change (FC) and p-value estimates for sirC versus nsirC and sirC versus HC comparisons are provided.

**Supplementary Figure S3. Systemic and colonic immune features in patients with irC. A.** Summary of treatment history and histopathologic assessment of colon biopsies from the biopsy cohort of patients with irC included in the analysis. **B.** Serum IL17A concentrations in patients with and without irC from the experimental cohort, stratified by checkpoint inhibitor regimen (anti-PD-1, anti-CTLA-4). Multiple unpaired two-tailed *t*-tests were used to analyze the data; significant differences were observed in the anti-PD-1 cohort (HC vs. nsirC, HC vs. sirC; $P < 0.05$, denoted by asterisks), while differences in the anti-CTLA-4 cohort did not reach statistical significance. **C.** Representative multiplex IHC images of FFPE colon biopsies from four irC patients spanning a range of colitis severity (ICI enterocolitis, moderately active colitis, severely active colitis). Sections were stained for CD4 (green), CD8 (yellow), CD11b (cyan), CD68a (red), Arg1b (white), and Foxp3 (magenta) (scale bar 100 µm). **D.** Quantification of relative cluster frequency (%) for $CD11b^{+}CD68^{-}$, $CD4^{+}$, $CD4^{+}Foxp3^{+}$, $CD8^{+}$, and $CD68b^{+}$ populations from multiplex IHC images across patients grouped by histologic severity (enterocolitis, mild, active, severe). **E.** Multiplex evaluation of human irC colon tissue using two antibody panels with digital cell maps aligning FcγR-expressing and non-expressing cell populations. Left: representative multiplex IHC images stained for CD4, CD8, CD11b, Foxp3, and CD68b (scale bar 100 µm), with corresponding digital cell phenotype map color-coded by FcγR and lineage marker combinations (e.g., FcγRI, FcγRIIb, FcγRIIIa, $CD68^{+}$, $CD11b^{+}$, $CD19^{+}FcγRIIb^{+}$, $CD68^{+}FcγRI/IIIa$). Right: serial sections stained by chromogenic IHC for CD64 (FcγRI), CD32b (FcγRIIb), and CD16 (FcγRIIIa) (top), with corresponding multiplex overlay images of CD64, CD32b, CD16, CD11b, and CD68a (bottom).

**Supplementary Figure S4. Modeling irC in hFcyR mice. A.** Body weight change from day 0 to day 21 in the indicated experimental groups. We monitored body weight longitudinally and compared groups. **B.** Representative H&E-stained intestinal (Swiss roll) sections from control C57BL/6 mice following transfer of nsirC- or sirC-derived patient IgG in combination with anti-PD-1 or anti-CTLA-4, showing no overt immune cell aggregation or histopathologic alteration regardless of IgG source or checkpoint inhibitor. **C.** Representative H&E-stained intestinal (Swiss roll) sections from hFcγR mice following transfer of nsirC- or sirC-derived patient IgG in combination with anti-PD-1 or anti-CTLA-4. sirC IgG was associated with mild intestinal inflammation characterized by increased lamina propria cellularity, mucosal hyperplasia, and submucosal leukocytic infiltration, in contrast to nsirC IgG. **D.** Representative H&E-stained colon sections from hFcγR mice (n = 3 per condition per patient) after transfer of IgG from three sirC patients (Patient 1, ID: 071; Patient 2, ID: 035; Patient 3, ID: 211), with or without anti-PD-1 treatment, to model the clinical setting. Patient-derived IgG alone did not appreciably alter colonic architecture, whereas combined IgG and anti-PD-1 treatment was associated with increased leukocyte infiltration.

**Supplementary Figure S5. Additional features of irC in hFcγR mice.**
**A.** Donor-level leukocyte infiltration analysis, in which histology scores were averaged across all recipient mice for each IgG donor (HC, nsirC, sirC; n=3-11 donors per group). The direction of effect is consistent with the mouse-level comparison in **Fig. 2C**, although statistical significance is not retained given the reduced number of independent donors. **B.** Serum ALT levels in hFcγR mice across the indicated experimental groups. **C.** Serum cytokine levels in anti-PD-1 treated mice, highlighting increased IL17A levels consistent with findings in human irC. **D.** Composition and relative abundance of the colonic bacterial community, assessed by 16S rRNA gene sequencing of fecal samples collected at day 0 (baseline, before serum IgG transfer), day 13 (during inflammation progression), and day 20 (peak leukocyte infiltration).

**Supplementary Figure S6. Histologic and multiplex immunohistochemical characterization of immune checkpoint colitis modeled in hFcγR mice. A.** Representative H&E-stained colon sections from hFcγR mice (n = 2–3 per patient-derived IgG condition) treated with anti–CTLA-4 or anti–PD-1. **B.** Bar plots quantifying immune cell populations across the indicated experimental conditions. **C.** Representative multiplex IHC images of formalin-fixed, paraffin-embedded (FFPE) colon tissue from hFcγR mice in each experimental group. Sections were stained for CD3 (orange), CD19 (blue), CD4 (green), CD8 (yellow), F4/80 (red), CD11b (cyan), and FOXP3 (magenta). **D.** Cell-to-cell distance analysis showing that CD11b+ myeloid cells are located farther from FOXP3+ regulatory T cells than CD4+ or CD8+ T cells in the mouse colon.

**Supplementary Figure S7. Single-cell RNA-seq profiling of colonic immune cells after ICC modeling. A.** Transfer of IgG from patients with GI toxicity mediates immune response in hFcyR mice but not in C57BL/6 mice. Bar plot showing no changes in the distribution of cell populations across clusters in the colons of control C57BL/6 mice, color-coded. **B.** Global distribution of major $CD45^{+}$ immune cell subsets in the lamina propria of hFcγR mice across all experimental groups. **C.** Feature plots showing expression of representative marker genes used to define transcriptionally distinct clusters.

**Supplementary Figure S8. Pro-inflammatory transcriptional profiles and global cell distribution across experimental conditions. A.** Dot plot analysis of cytotoxic and cytokine/effector gene expression (*Gzma, Gzmb, Gzmc, Ifng, Il10, Il13, Il17a, Il1b, Il2, Il22, Il4, Il5, Il6, Prf1*) across immune cell clusters identified by scRNA-seq (Mac, MON, DC, B-cycling, B, PC, NK+ILC1, ILC2, ILC3, Tgd, CD8, CD4, ITLeff, ITL), shown for all datasets combined (top) and separately for sirC IgG + anti-PD-1 (middle) and sirC IgG + anti-CTLA-4 (bottom) conditions. Dot size and color represent relative expression level, row-normalized from row minimum (blue) to row maximum (red). **B.** UMAP projections of $CD45^{+}$ lamina propria cells, colored by cluster identity, shown separately for each of the eight experimental conditions (HC IgG + anti-CTLA-4, HC IgG + anti-PD-1, nsirC IgG + anti-CTLA-4, nsirC IgG + anti-PD-1, sirC IgG + anti-CTLA-4, sirC IgG + anti-PD-1, sirC IgG + anti-CTLA-4 + isotype control, sirC IgG + anti-PD-1 + isotype control), illustrating differences in overall immune cell composition and distribution across treatment groups. **C.** Principal component analysis (PCA) comparing global transcriptional profiles across all experimental conditions within the anti-PD-1 (orange circles) and anti-CTLA-4 (blue triangles) treatment arms. PC1 and PC2 account for 18.5% and 71.0% of variance, respectively. HC, healthy control; NT, nsirC (non-severe irC); T, sirC (severe irC); Iso, isotype control.

**Supplementary Figure S1.**

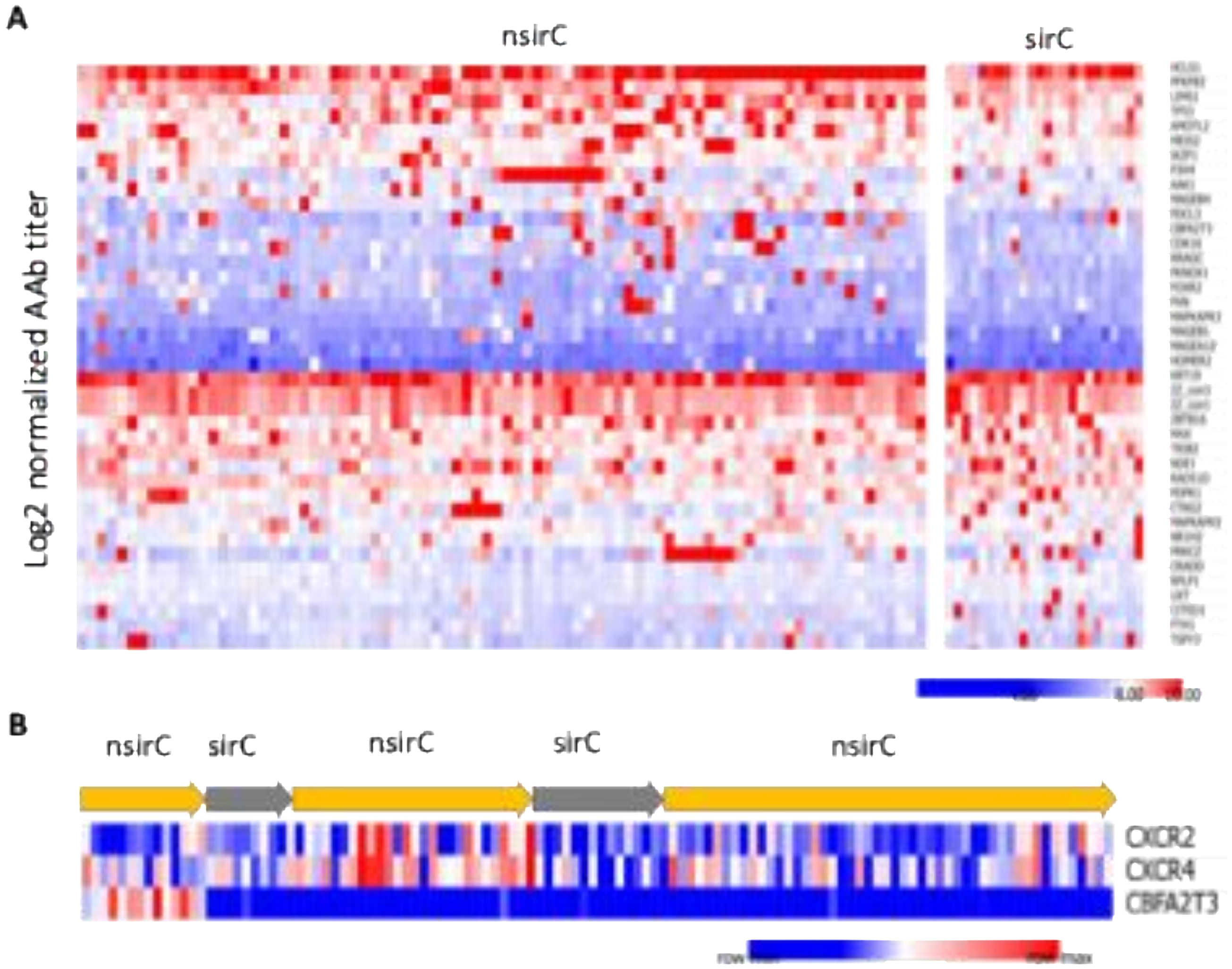

**Supplementary Figure S2**.

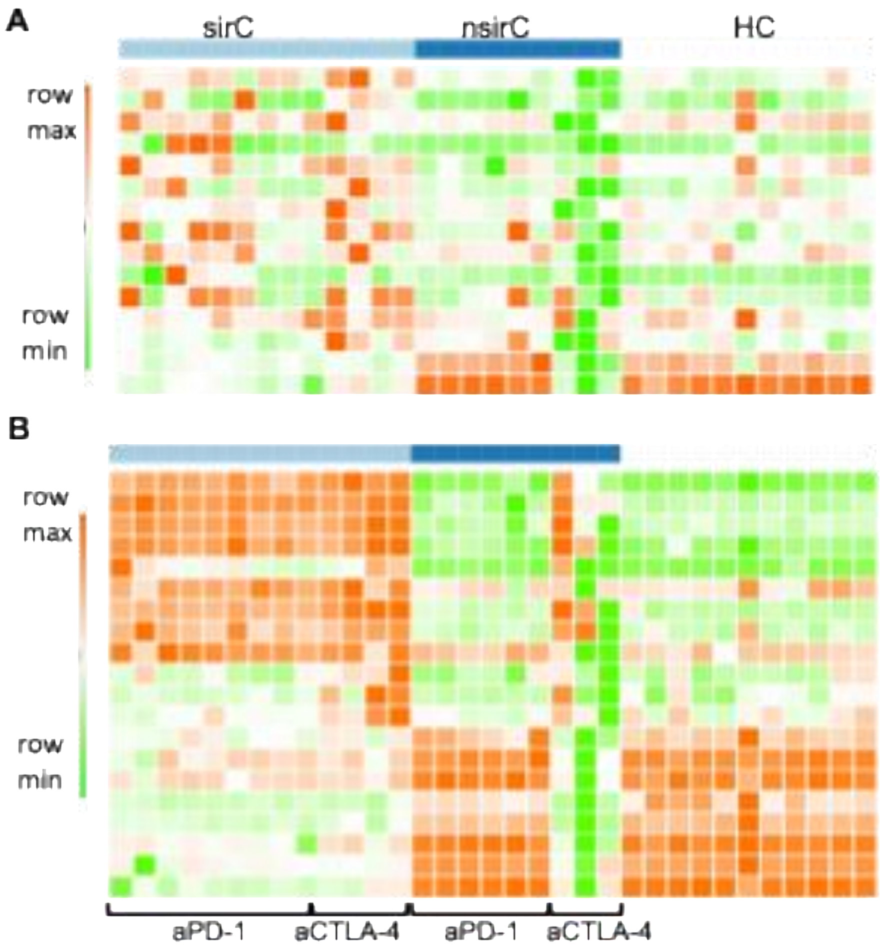


| | sirC vs nsirC | | sirC vs HC | |
|---|---|---|---|---|
| | FC | p_value | FC | p_value |
| AKAP4 | 3.19 | 0.01 | 3.88 | 0.01 |
| MLANA | 3.69 | 0.01 | 0.95 | 0.35 |
| CT47A1 | 2.66 | 0.02 | 0.36 | 0.72 |
| GAGE1 | 2.47 | 0.03 | 1.45 | 0.17 |
| TMEFF2 | 2.46 | 0.03 | 0.96 | 0.35 |
| MAGEA10 | 2.27 | 0.04 | 1.32 | 0.2 |
| EWSR1 | 2.17 | 0.05 | 0.83 | 0.42 |
| CTAG2 | 1.91 | 0.07 | 3.99 | 0.01 |
| PMEL | 1.86 | 0.08 | 0.91 | 0.37 |
| MAGEA4 | 1.63 | 0.12 | 2.12 | 0.05 |
| CTAG1A | 1.55 | 0.14 | 2.72 | 0.02 |
| ENO1 | 1.35 | 0.2 | 0.58 | 0.57 |
| BRAF | 0.58 | 0.57 | 1.27 | 0.22 |
| BRD2 | -1.29 | 0.23 | -6.41 | 0 |
| BASP1 | -2.16 | 0.06 | -12.34 | 0 |

| | FC | p_value | FC | p_value |
|---|---|---|---|---|
| EEF1D | 6.4 | 0 | 26.29 | 0.00 |
| STK17B | 5.41 | 0 | 22.43 | 0.00 |
| PDK4 | 5.05 | 0 | 19.21 | 0.00 |
| HES6 | 4.79 | 0 | 15.68 | 0.00 |
| GSTT1 | 3.91 | 0 | 7.65 | 0.00 |
| CDK5R1 | 3.87 | 0 | 3.46 | 0.00 |
| DMRTB1 | 3.68 | 0 | 12.92 | 0.00 |
| PRKAA2 | 3.33 | 0.01 | 7.03 | 0.00 |
| STK4 | 2.8 | 0.02 | 5.23 | 0.00 |
| FADD | 2.38 | 0.03 | 1.57 | 0.13 |
| MX1 | 1.7 | 0.11 | 1.39 | 0.18 |
| MAX | 1.56 | 0.14 | 0.78 | 0.44 |
| PRKACA | -0.64 | 0.54 | -5.30 | 0.00 |
| FGFR1 | -1.16 | 0.28 | -10.23 | 0.00 |
| FKBP3 | -1.24 | 0.25 | -10.67 | 0.00 |
| C6 | -1.28 | 0.23 | -6.90 | 0.00 |
| HTR1D | -1.47 | 0.18 | -12.04 | 0.00 |
| HRH2 | -1.65 | 0.13 | -12.11 | 0.00 |
| OVOL1 | -2.42 | 0.03 | -8.68 | 0.00 |
| TCF12 | -3.19 | 0.01 | -15.56 | 0.00 |

**Supplementary Figure S3.**

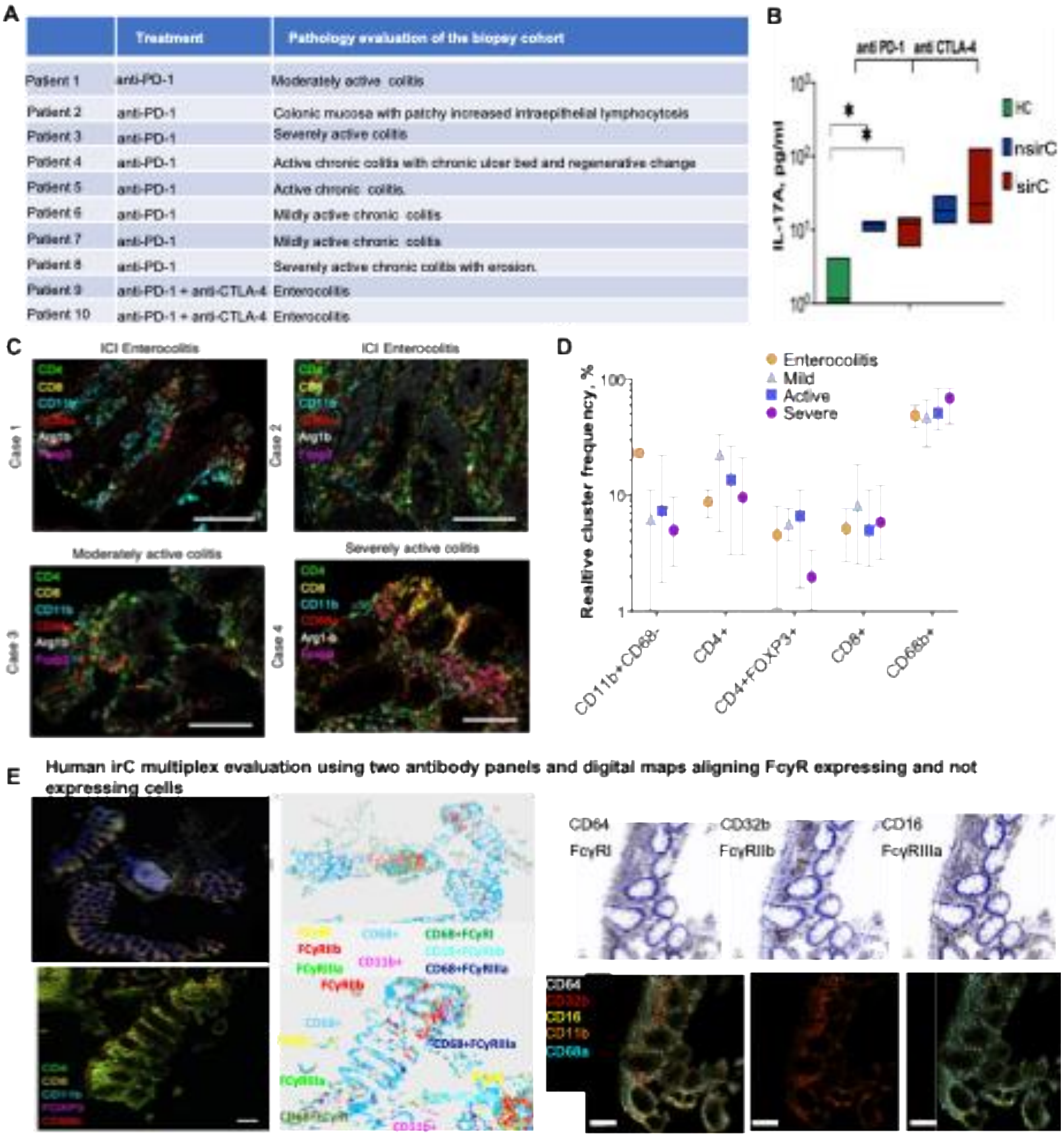

| | Treatment | Pathology evaluation of the biopsy cohort |
|---|---|---|
| Patient 1 | anti-PD-1 | Moderately active colitis |
| Patient 2 | anti-PD-1 | Colonic mucosa with patchy increased intraepithelial lymphocytosis |
| Patient 3 | anti-PD-1 | Severely active colitis |
| Patient 4 | anti-PD-1 | Active chronic colitis with chronic ulcer bed and regenerative change |
| Patient 5 | anti-PD-1 | Active chronic colitis. |
| Patient 6 | anti-PD-1 | Mildly active chronic colitis |
| Patient 7 | anti-PD-1 | Mildly active chronic colitis |
| Patient 8 | anti-PD-1 | Severely active chronic colitis with erosion. |
| Patient 9 | anti-PD-1 + anti-CTLA-4 | Enterocolitis |
| Patient 10 | anti-PD-1 + anti-CTLA-4 | Enterocolitis |

**Supplementary Figure S4.**

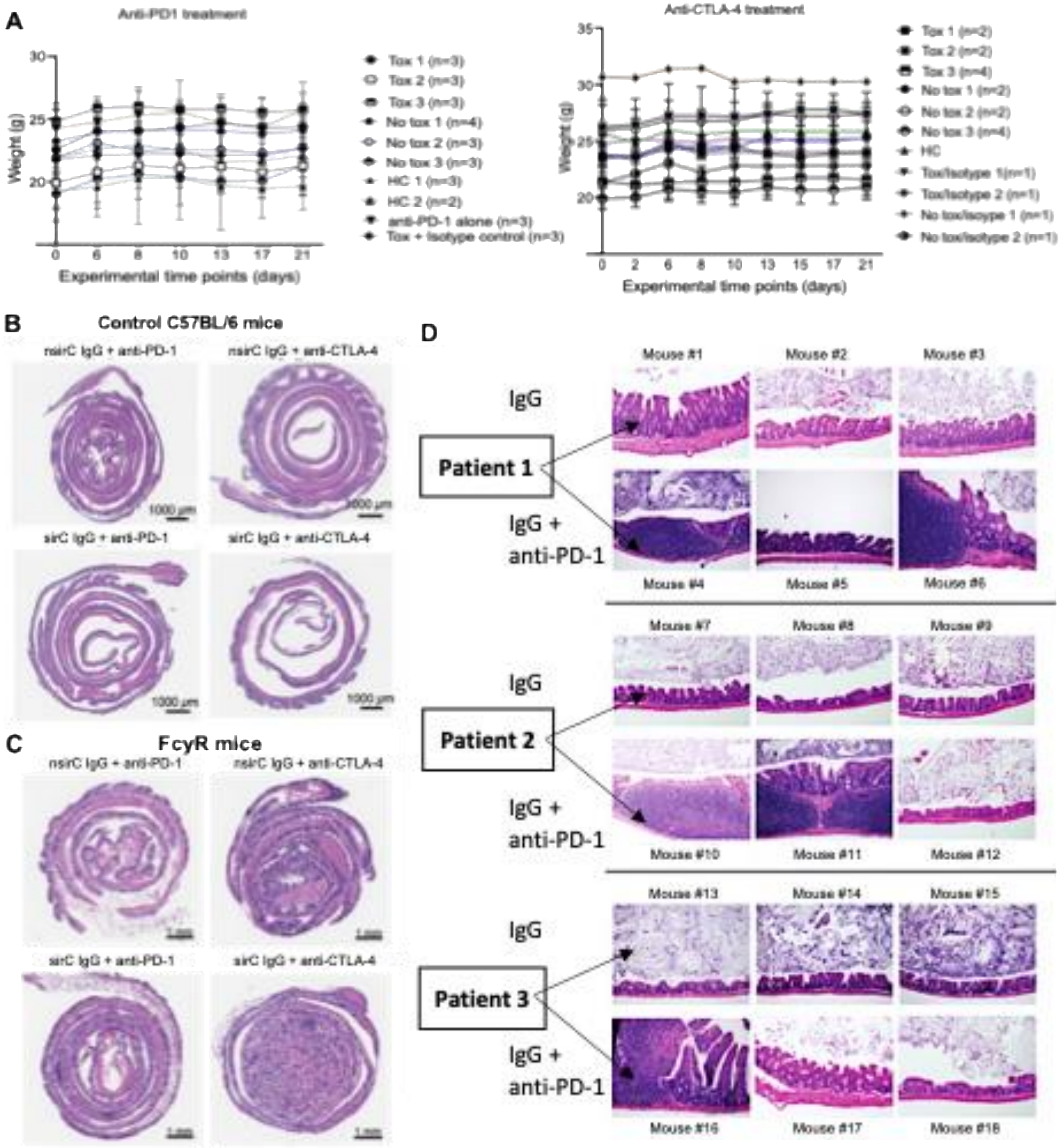

**Supplementary Figure S5.**

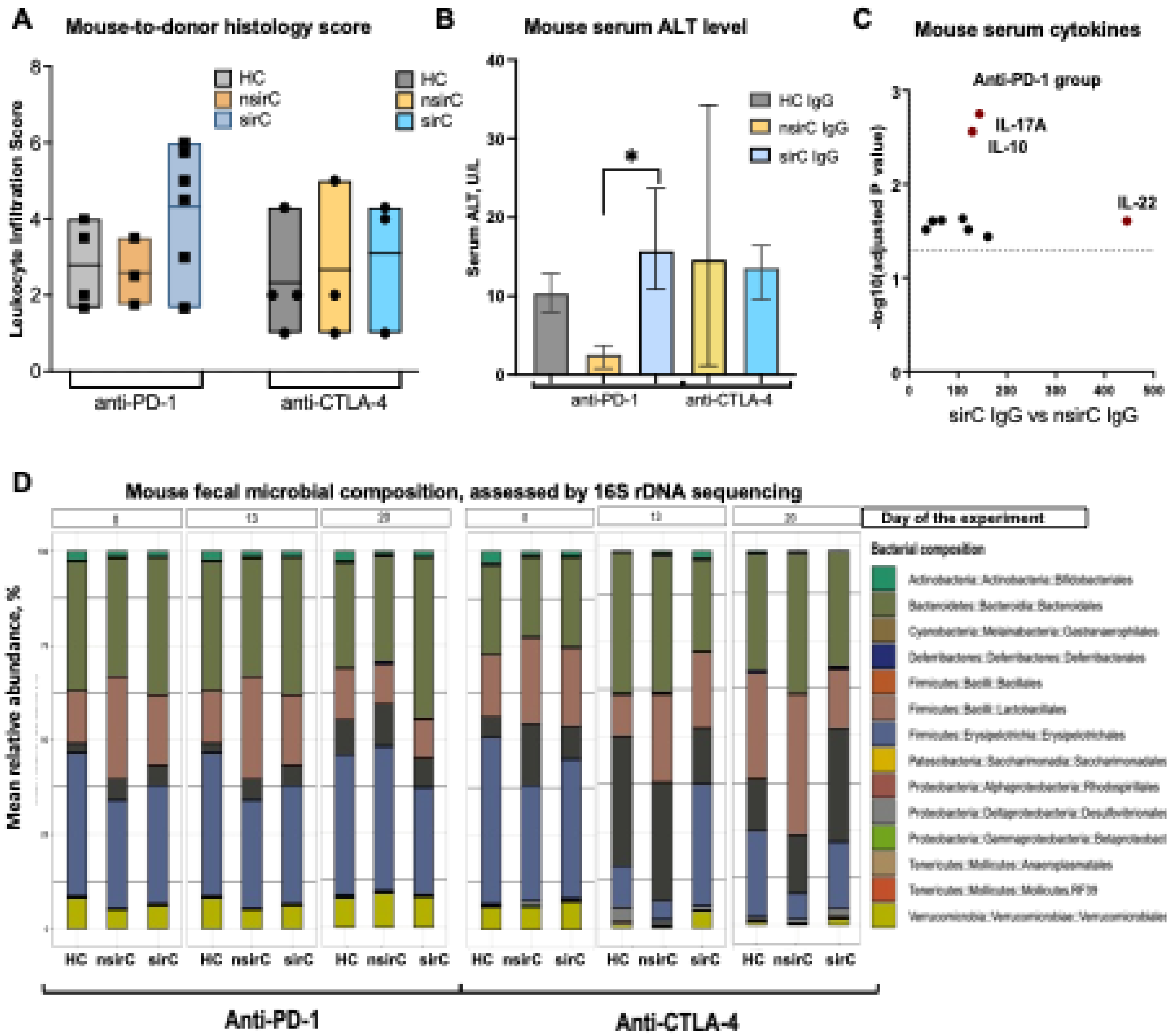

**Supplementary Figure S6.**

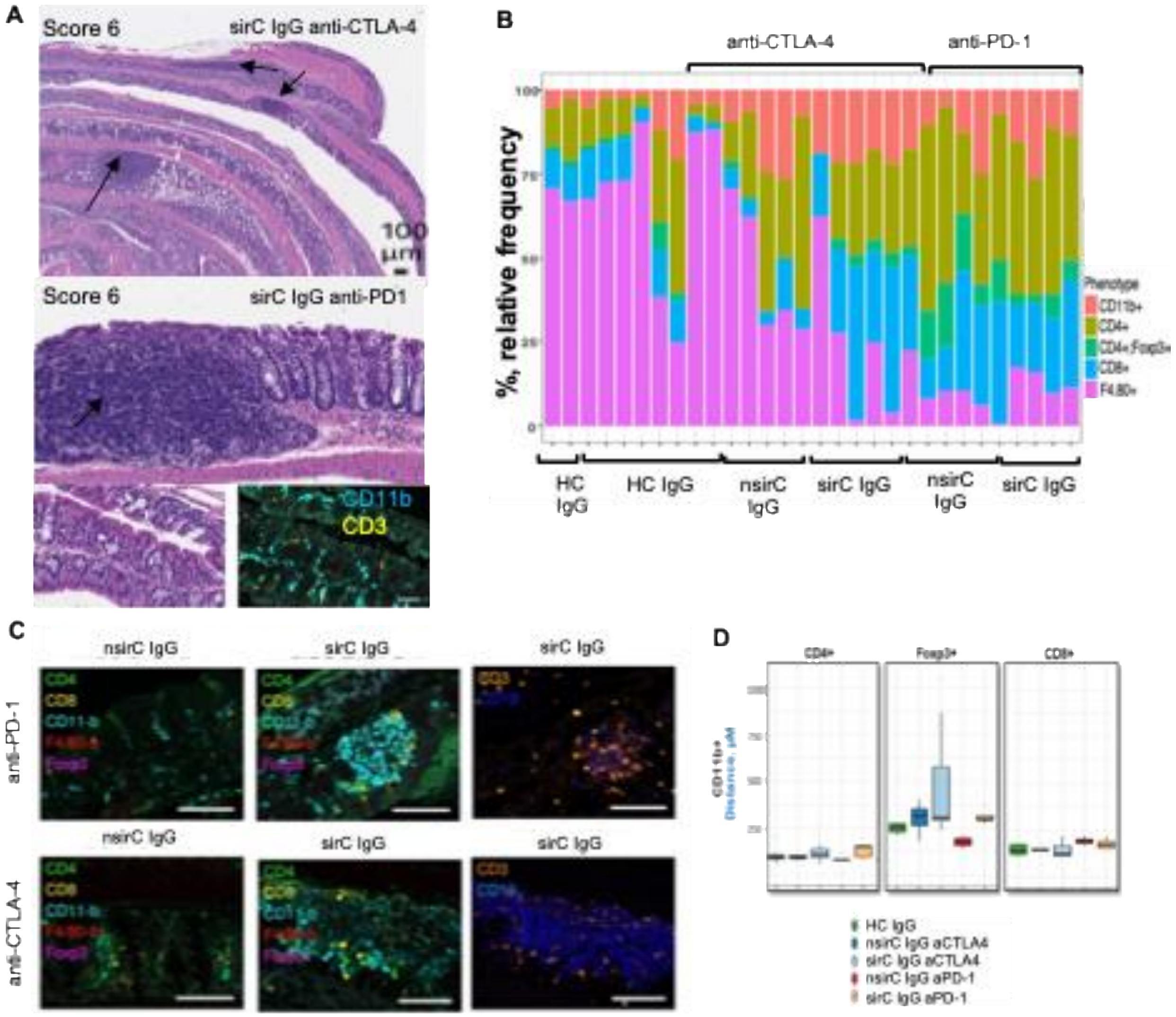

**Supplementary Figure S7.**

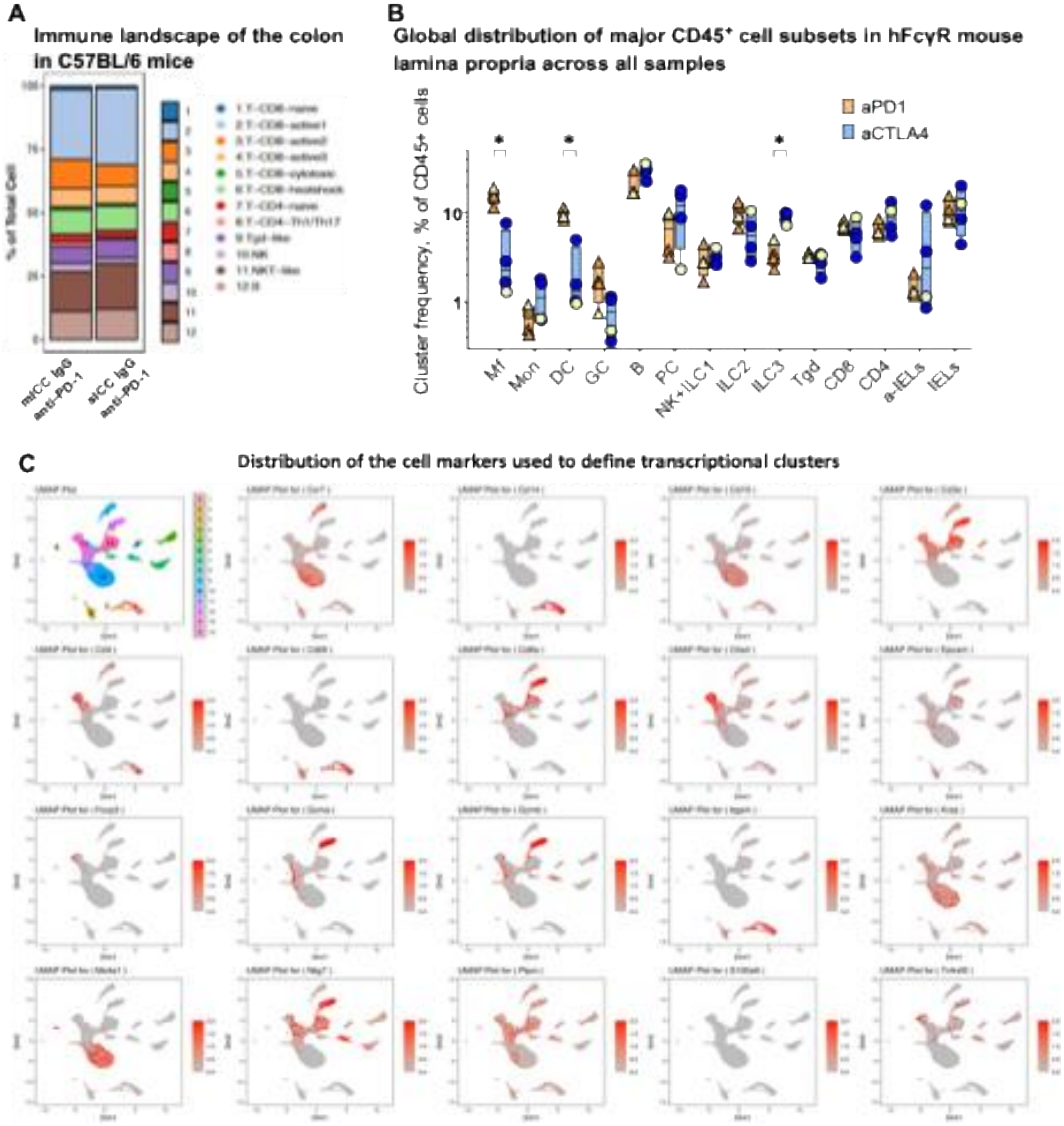

**Supplementary Figure S8.**

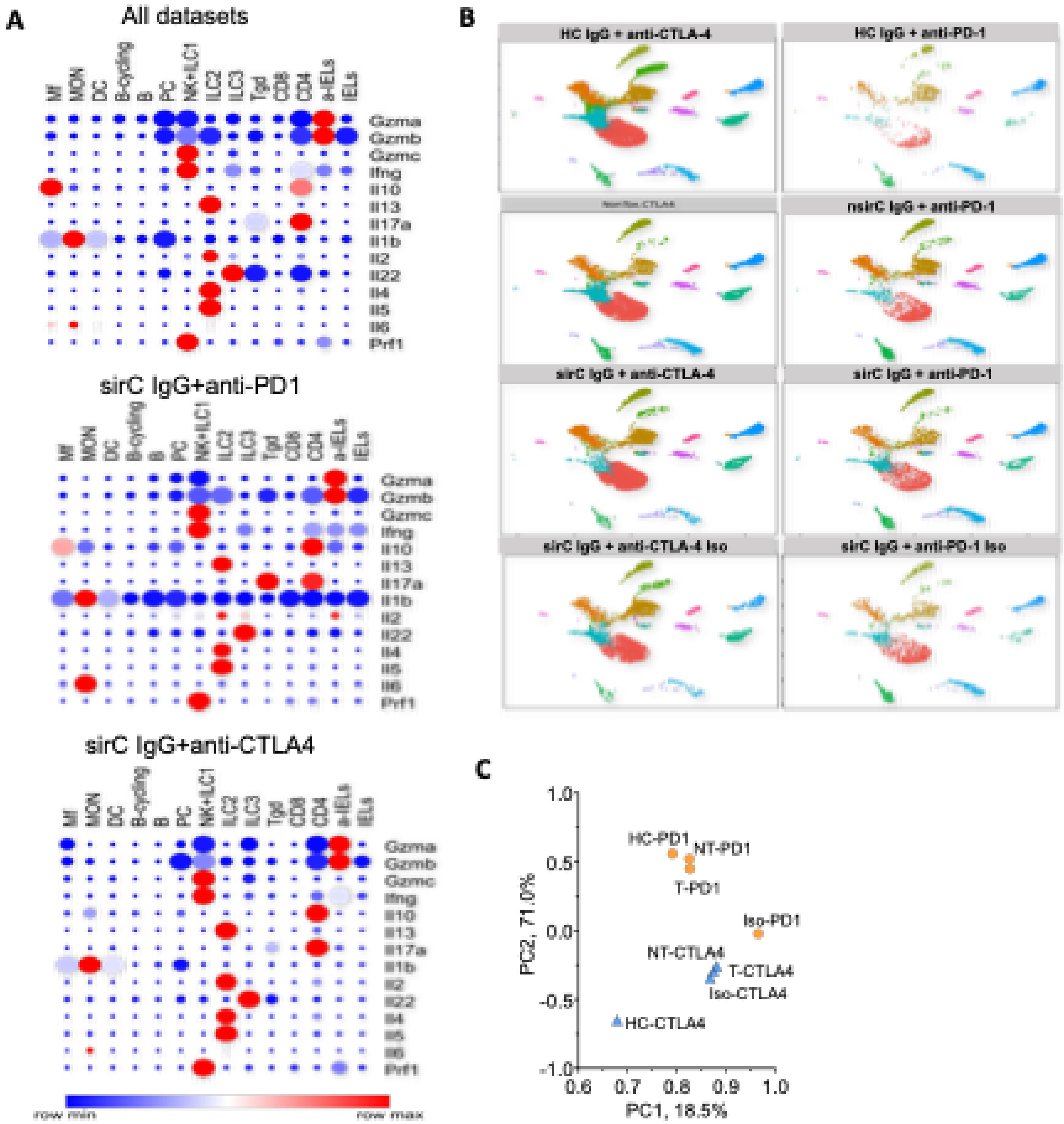
